# The Piezo channel is used by migrating cells to sense compressive load

**DOI:** 10.1101/534172

**Authors:** Nishit Srivastava, David Traynor, Alexandre J. Kabla, Robert R. Kay

## Abstract

Migrating cells face varied mechanical and physical barriers in physiological environments, but how they sense and respond to them remains to be fully understood. We used a custom-built ‘cell squasher’ to apply uniaxial pressure to *Dictyostelium* cells migrating under soft agarose. Within 10 seconds of application, loads of as little as 100 Pa cause cells to move using blebs instead of pseudopods. Cells lose volume and surface area under pressure and their actin dynamics are perturbed. Myosin-II is recruited to the cortex, potentially increasing contractility and so driving blebbing. The blebbing response depends on extra-cellular calcium, is accompanied by increased cytosolic calcium and largely abrogated in null mutants of the Piezo stretch-operated channel. We propose that migrating cells sense mechanical force through mechano-sensitive channels, leading to an influx of calcium and cortical recruitment of myosin, thus re-directing the motile apparatus to produce blebs rather than pseudopods.

## Introduction

Cell movement is a key way in which animals shape their body during embryonic development, and defend and repair it as adults ^1–4^. In the body, motile cells have to navigate through complex three-dimensional environments in which they can encounter various mechanical challenges, such as obstacles, narrow spaces, barrier membranes and resistance from the extracellular matrix ^5,6^.

To overcome these obstacles cells need both sensors for the physical properties of the environment, allowing them to detect the forces that it applies to them, and the ability to adjust their movement accordingly ^7^. For instance, a cell moving through packed tissue may require much greater force at the leading edge than one under buffer, while greater contractile forces may be needed to draw the cell body and nucleus forward through constricted spaces and against friction from the surrounding tissue ^8,9^. Similarly, cells need to avoid absolute barriers, and it might be advantageous if they could select paths offering the least mechanical resistance ^10–12^.

Cells can respond aggressively to mechanical barriers like the extra-cellular matrix by using specialist organelles such as podosomes to degrade them with secreted enzymes ^13,14^. In many cases cells also change the way in which they move to give greater emphasis to contractile forces driven by acto-myosin contraction ^15^. These forces produce a pressurised cytoplasm that can help to drive forward the leading edge, resulting in the formation of blebs in the extreme case. Such bleb-driven movement was described more than 40 years ago in fish embryos, and a greater reliance on myosin contractility is a common feature of cells moving in 3-dimensional environments, including tumour cells ^16–19^.

Mechanical force can be sensed in a variety of ways. The branched F-actin network at the leading edge of migrating cells is force-sensitive and intrinsically adapts to load by increasing the density of F-actin filaments and changing their angle to the plasma membrane ^20,21^. Cells can also sense mechanical force using stretchable proteins, such as talin, as strain gauges ^22–24^ and the actin cytoskeleton itself is responsive to strain ^25^. Stretch-operated channels in the plasma membrane are also used to sense mechanical forces in both prokaryotes and eukaryotes ^26,27^. Most relevant here is the Piezo channel, which can be opened by strain in the membrane and lets through a variety of cations, including calcium ^28–30^; it is responsible for light touch sensation and sensing of crowding in epithelia among many other things ^31–33^, and is a strong candidate for mechanical sensing in cell migration^34^.

The very complexity of natural cellular environments can make it hard to tease out the contribution of individual mechanical and chemical influences on cell behaviour ^35^. It is often unclear exactly what features cells are responding to, nor the sensing system mediating their response. Simplified systems are therefore useful to analyse this complexity.

Like animal cells, *Dictyostelium* amoebae move through varied environments during their life cycle. In their single-celled growth phase, they move through the interstices of the soil hunting for bacteria, while in their developmental stages they form multicellular aggregates where cell movement and cell sorting within the aggregate play important roles in morphogenesis ^36,37^. *Dictyostelium* cells can extend their leading edge using pseudopods driven by actin polymerization, or blebs driven by fluid pressure and dependent on myosin-II ^38–40^. We found previously that cells prefer pseudopods when moving under buffer, but blebs under a stiff agarose overlay ^41^. In both cases the cells move on the same glass substratum, but under agarose they must also make space for themselves at the interface by breaking the adhesive forces between the substratum and the overlay and by deforming the overlay itself. The cells therefore experience both increased mechanical resistance at the leading edge and the elastic forces from the overlay compressing the cell body. It seems likely that one or both of these somehow trigger the switch to bleb-driven movement.

In order to study how mechanical forces trigger a change in movement mechanics we built a ‘cell squasher’ to rapidly apply defined loads to cells under an agarose overlay ^42^ while leaving other potential variables, such as chemical composition and degree of cross-linking of the matrix, or even oxygen availability, largely constant. Using *Dictyostelium* cells, this has allowed us to investigate one variable – the uni-axial load on cells – in isolation. We find that modest loads rapidly cause cells to switch to a bleb-driven mode of movement and that this depends almost entirely on the Piezo stretch-operated channel, most likely acting through a calcium signal to re-configure the motile apparatus towards myosin-II driven contractility.

## Results

### Uni-axial loads cause cells to rapidly adopt bleb-driven motility

We used a custom built ‘cell-squasher’ to apply uniaxial loads to *Dictyostelium* cells moving under a thin layer of agarose on a glass surface^42^ (Figure 1A). In most experiments, we used cells chemotaxing to cyclic-AMP and transformed with fluorescent reporters, so that they could be observed by confocal microscopy. Blebs and pseudopods were distinguished by their characteristic morphologies and dynamics, as revealed by an F-actin reporter ^41^ (Figure 1B). Blebs are rounded, expand abruptly and leave behind a characteristic F-actin scar (the former cell cortex), which dissipates over a few tens of seconds. Their membrane is initially free of F-actin, but rapidly reforms an F-actin cortex. In contrast, pseudopods are more irregular, expand steadily but more slowly, and always have F-actin at their leading edge.

**Figure 1.**
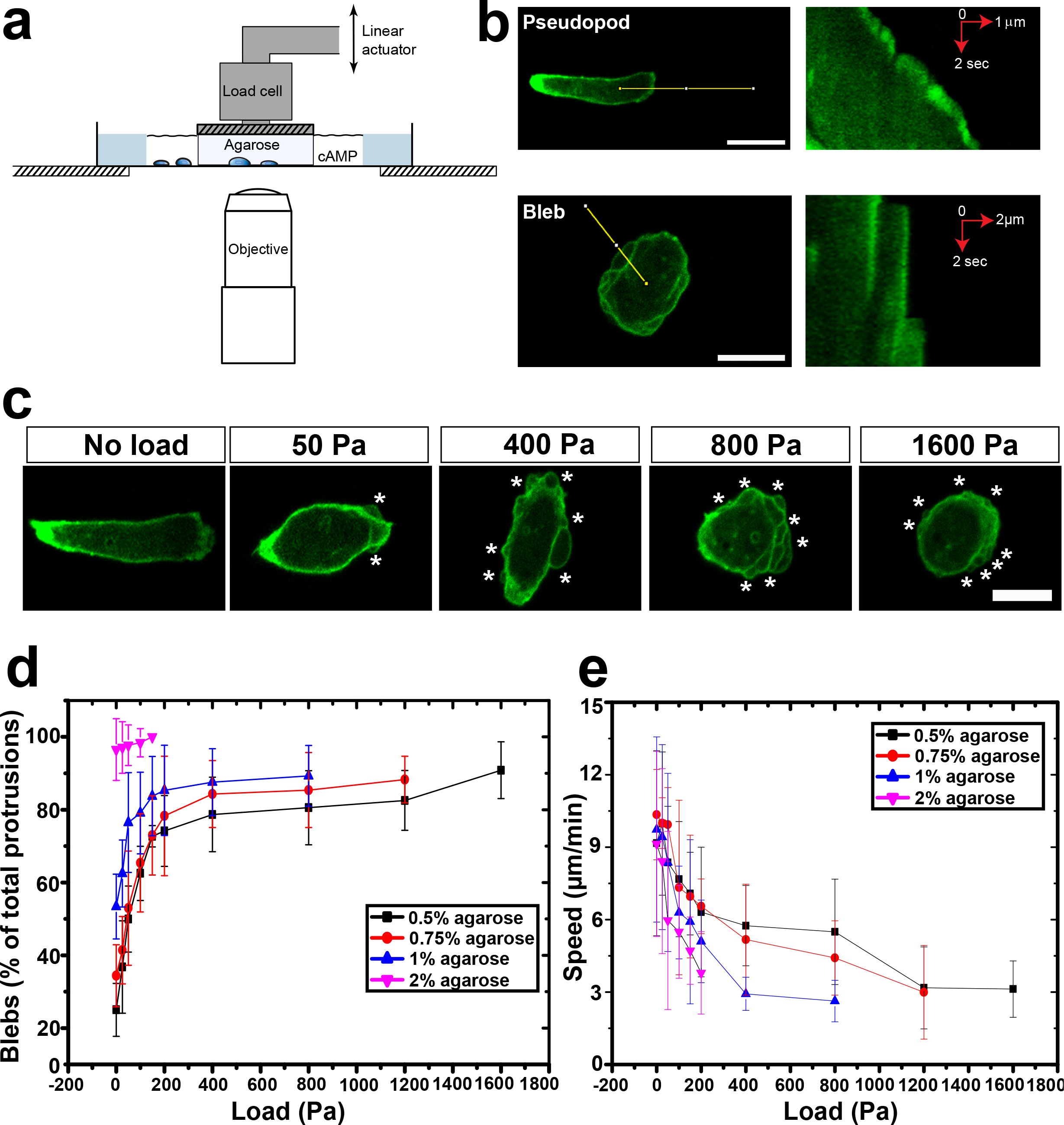
Uniaxial load causes cells to move using blebs instead of pseudopods. (A) Diagram of the ‘cell squasher’ used to apply uniaxial loads to cells moving under an agarose overlay^42^. The load is applied using a plunger on the bridge between two wells cut into the agarose, one containing cells and the other the chemoattractant cyclic-AMP, which attracts the cells under the agarose towards it. (B) Distinction between pseudopods and blebs. Left panels show cells expressing an F-actin reporter and the right panels, kymographs taken along the lines indicated in the left panels; scale bar = 10 μm. (C) Uniaxial load causes migrating cells to bleb. The cells are migrating under an overlay of 0.5% agarose to which increasing uniaxial loads are applied. Blebs are indicated by white stars; scale bar = 10 μm. (D) Blebbing of migrating cells increases with increasing load or overlay stiffness. (E) Cell speed decreases under increasing load or stiffness of the agarose overlay. The data is represented as mean ± SD for n ≥ 30 cells for each case with measurements made for about 30 minutes, starting 8-10 min after load was applied. The stiffness of the agarose overlays is: 0.5% = 6.6 kPa; 0.75% = 11.9 kPa; 1% = 20.5 kPa; 2% = 73.6 kPa. Aggregation-competent Ax2 cells expressing the ABD120-GFP reporter for F-actin and migrating towards cyclic-AMP are used throughout.

Cells under a soft 0.5% agarose overlay move using a mixture of both pseudopods and blebs, with pseudopods predominating (~75% of total large projections) (Figure 1C and D; movie S1). Their response to pressure is striking: blebbing is stimulated by pressures of as little as 50 Pa and is half-maximal at 100 Pa (Figure 1C and D; movie S2). The basal level of blebbing is greater at higher percentages of agarose in the overlay, but load again causes an increase in blebbing, which approaches 100% with cells under 2% agarose (Figure 1D). Cells also move more slowly under load, again proportional to the load applied and the stiffness of the overlay (movie S3 & S4).

The onset of blebbing is very rapid. Within 10 seconds of starting to apply load there is a clear increase in blebbing from about 2 to 10 blebs/cell/minute (Figure 2, movie S5). This may even be an underestimate of how quickly cells can respond, since with our device it is necessary to ramp up the load over 20 seconds to avoid a loss of focus. The increased rate of blebbing is sustained as long as the load is maintained but can be reversed in 8-10 minutes if it is removed, with cells again moving predominantly with pseudopods (Supplementary figure 1, movie S6). The quantification of protrusions confirms the trend with the cells forming only about 3 blebs/cell/min after the load is removed (Supplementary figure 1B).

**Figure 2.**
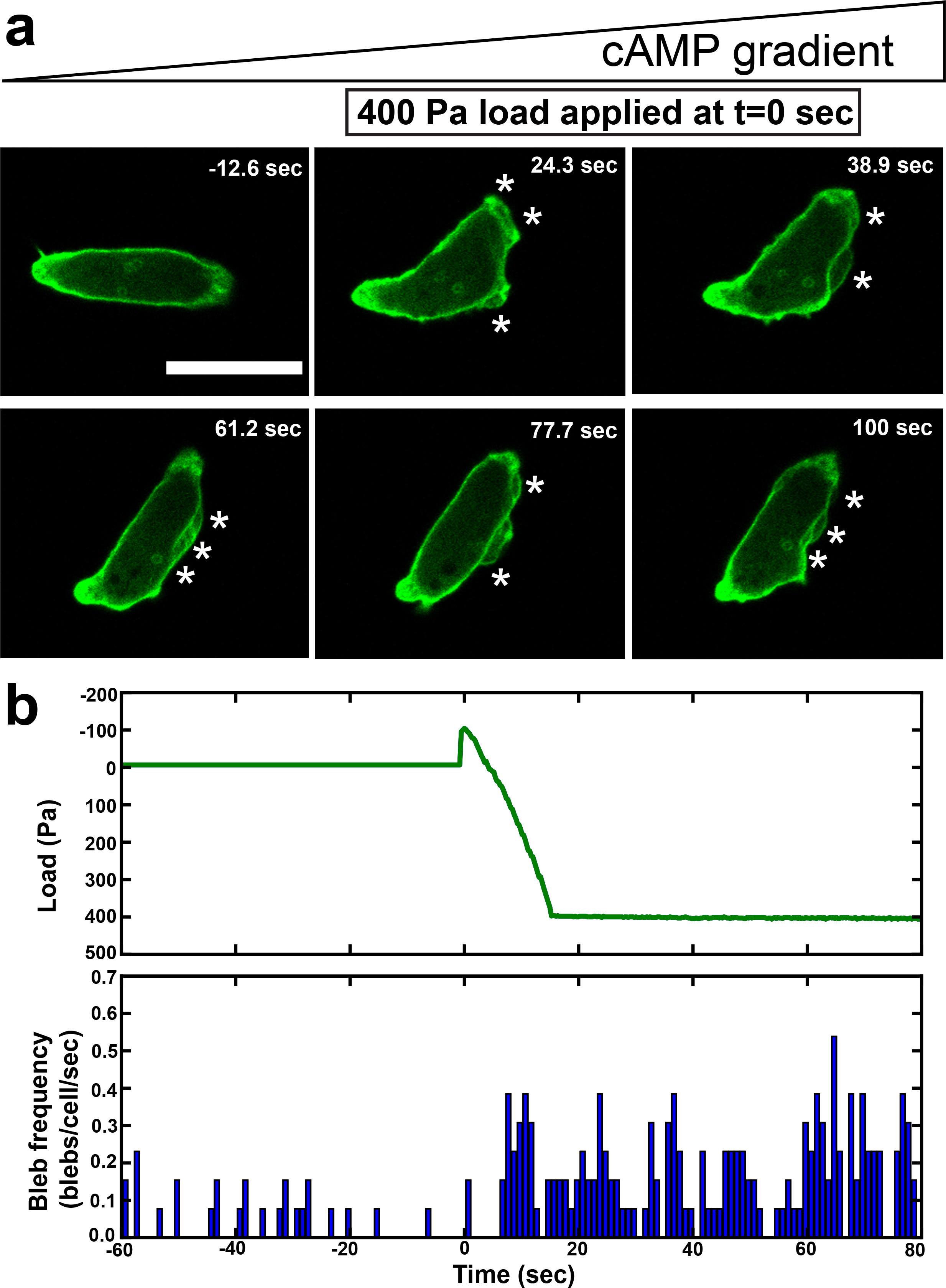
Uniaxial load causes a rapid switch to bleb-driven migration. (A) Rapid induction of blebbing by uniaxial loading of migrating cells. Frames from a movie timed with respect to the start of loading (t=0); blebs are indicated by * and scale bar =10 μm. (B) Time course of bleb induction by load. The top panel shows a typical loading regime with a small up-tick as the plunger first touches the agarose followed by a 15 sec ramping of load to 400 Pa. Bottom panel shows the bleb frequency, with blebs binned into 1 sec intervals and scored at the time they first appear (typically they are fully expanded in one frame of the movie). Aggregation-competent Ax2 cells expressing the F-actin reporter ABD120-GFP were filmed at 2 frames/sec under a 0.5% agarose overlay (N=17 cells).

These results demonstrate that uniaxial load alone is sufficient to dramatically change how cells move, directing them to use blebs instead of pseudopods, and the speed of response shows that cells must possess a fast-acting response system to mechanical load.

### Load causes cells to shrink and affects actin dynamics

To begin to understand how cells respond to load, we investigated its effects on cell morphology and the actin cytoskeleton (Table 1). Cells flatten under load, with a reduction in their height and volume as measured from 3D reconstructions. For instance, a load of 400 Pa applied to cells under 0.5% agarose, causes their height to decrease from 7 ± 1 μm to 3 ± 1 μm and volume by 25% (480 ± 70 μm^3^ to 360 ± 60 μm^3^).

**Table1:**
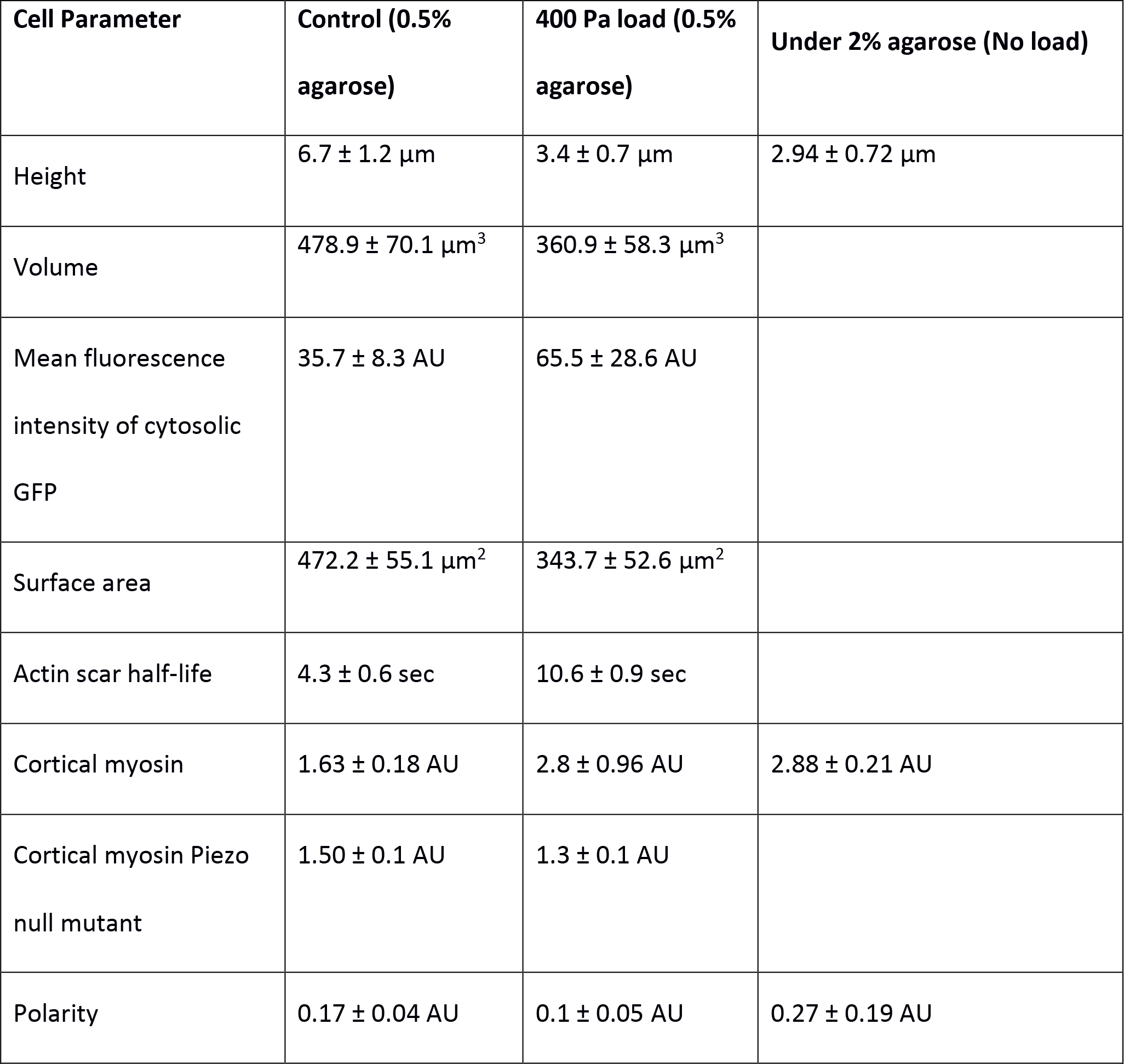
The effect of load and overlay stiffness on cellular parameters. The effect of load or overlay stiffness on cellular parameters was determined in the conditions shown. Aggregation-competent cells migrating towards cyclic-AMP were used in all cases with either 0.5% or 2% agarose overlays, giving stiffness of 6.6 kPa and 73.6 Pa respectively. Ax2 cells were used, except where indicated (n>30 in all cases). The methods used to quantify cellular parameters are described in the Materials and Methods.

The surface area of freely moving *Dictyostelium* cells can change by as much as 30% over a few minutes, but volume tends to be much more stable ^43^, and we were therefore surprised to find that cell volume changes significantly under load. We used measurements of the intensity of soluble GFP in confocal sections as an independent way to detect changes in cytoplasmic volume (Table 1). In this case, since GFP is not lost from cells when they shrink, an increase in fluorescence indicates an increased GFP concentration and a loss of cell volume. This approach reports an even larger decrease in volume of around 44% (fluorescence increased from 35.7 ± 8.3 AU to 65.5 ± 28.6 AU under 400 Pa load and 0.5% agarose gel (Table 1). The discrepancy between methods might be explained if the volume loss occurs only from the cytoplasm, where GFP resides, and not from other organelles such as vesicles.

Compressive load also causes a loss of polarity. Without load, cells are typically polarized with a pronounced leading edge directed up the cyclic-AMP gradient (Figure 1C). At loads above 100 Pa, the cells become less polarized, concomitant with the appearance of blebs. At higher load still, the cells round up with a complete loss of a distinct leading-edge (Figure 1C).

The actin cytoskeleton is substantially perturbed under load. Coronin ^44,45^, an F-actin binding protein required for efficient chemotaxis, redistributes from pseudopods to the F-actin scars left by expanding blebs (Figure 3A), and these scars tend to linger for a much longer time. Scar lifetime was quantitated by fitting a sigmoidal to the decay of fluorescence of an F-actin reporter in the scar (Supplementary figure 2 and Material and Methods) and found to increase from a half-life of 4 ± 1 seconds in control cells to 11 ± 1 under 400 Pa and 13 ± 7 seconds under 800 Pa load, so that the cell perimeter becomes marked by slowly-dissipating arcs of F-actin (Table 1).

**Figure 3.**
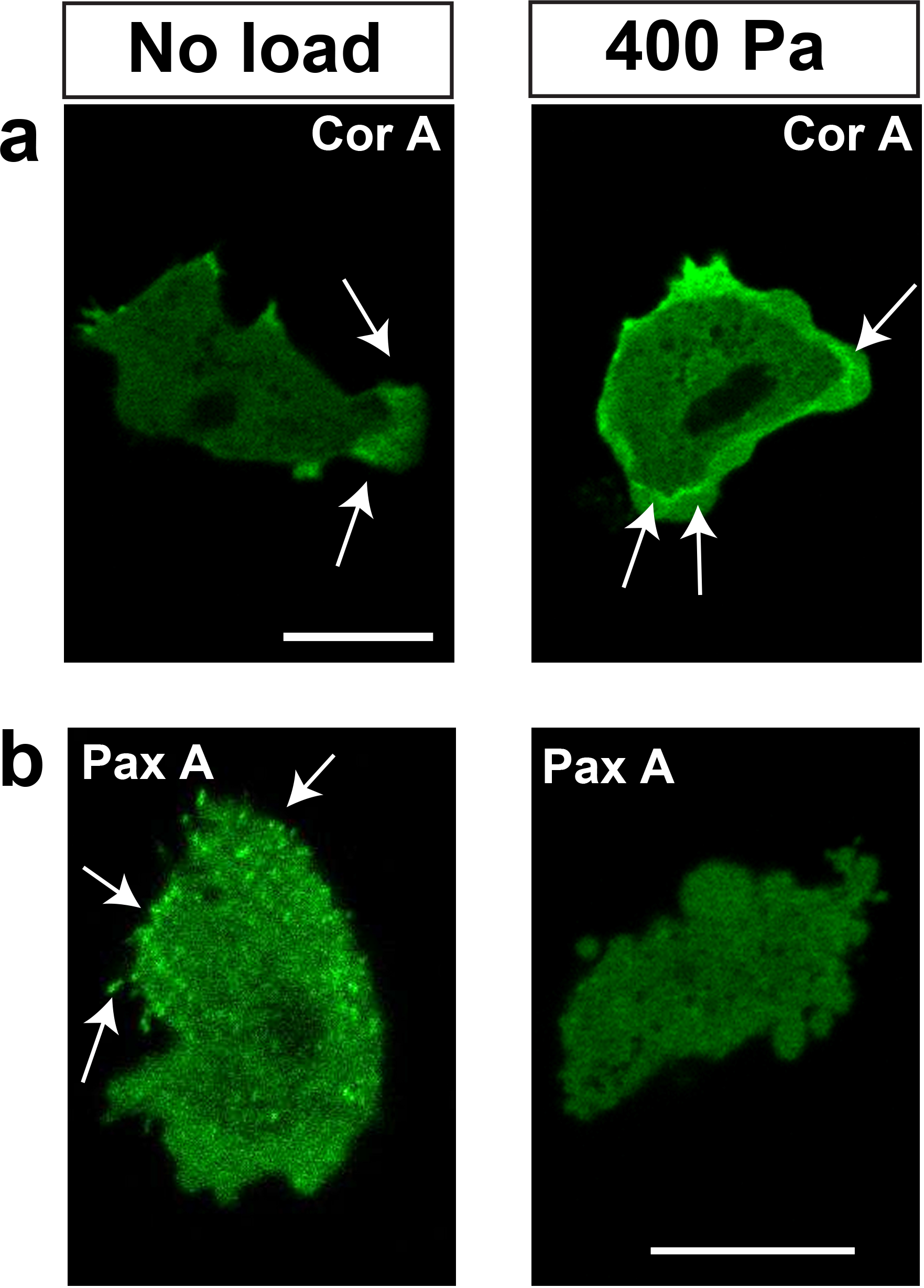
Uniaxial load causes cytoskeletal re-organization. (A) Coronin, an F-actin binding protein, relocates under load from pseudopods to the actin scars left behind by blebs. (B) Paxillin patches, thought to mediate adhesion to the substratum, disperse under load. Load was applied to aggregation-competent Ax2 cells expressing either the coronin-GFP reporter or GFP-paxillin and migrating towards cyclic-AMP, under an overlay of 0.5% agarose; scale bar = 10 μm.

The basal surface of migrating cells is decorated by punctate local adhesions containing paxillin that can be visualised with paxillin-GFP (Figure 3B) ^46^. These tend to dissipate under load suggesting that cells become less adhesive when moving with blebs (Figure 3B).

### Load causes myosin-II recruitment to the cell cortex

We next tried to trace the steps linking applied load to increased blebbing. Blebs are driven out by fluid pressure and since the cytoplasm is pressurised by contraction of the cell cortex, which depends on myosin-II ^39,41^, we reasoned that myosin-II might be recruited and activated at the cell cortex. In accord with this idea, myosin-II null cells ^47^, which are incapable of blebbing in normal physiological circumstances ^39,41^ (Supplementary figure S4D) do not bleb when load is applied. This shows that blebbing in response to load depends on myosin-II and is likely an active response of cells to load.

A GFP-myosin-II reporter expressed in myosin-II null cells restores their ability to bleb and is therefore functional. This reporter accumulates preferentially in the cortex towards the rear of control cells chemotaxing with pseudopods as their primary protrusions; however a 400 Pa load causes a sudden and more uniform recruitment to the cortex (Figure 4A, movie S7), with the cortical enrichment increasing from by about 60% (Figure 4B, Supplementary figure S3). Recruitment of myosin-II is rapid, occurring in less than 20 seconds and so is on the same scale as the increase in blebbing (Figure 4C).

**Figure 4.**
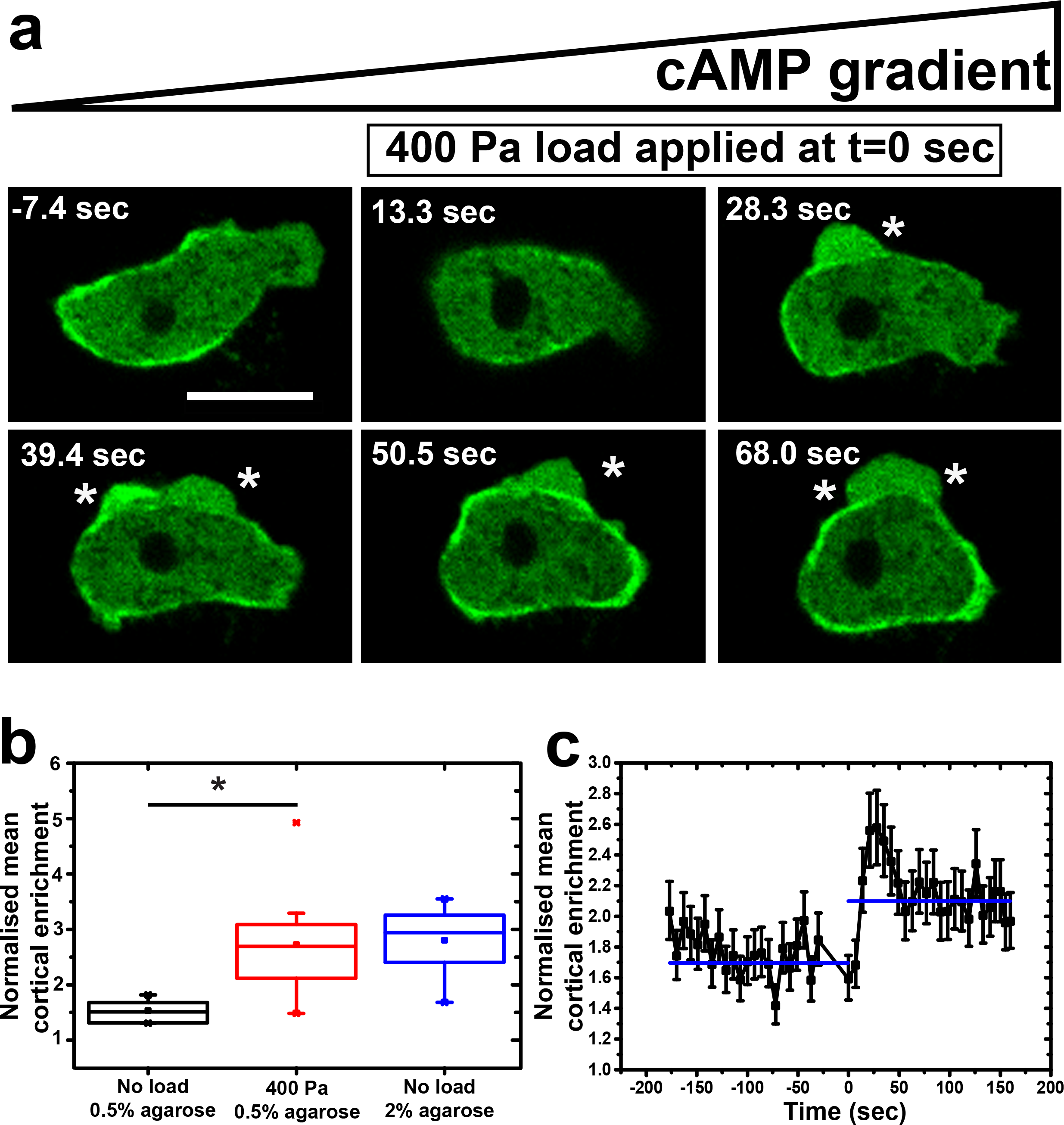
Myosin-II is rapidly recruited to the cell cortex in response to load. (A) Load causes myosin-II to be recruited to the cell cortex. Blebs are indicated by * and scale bar = 10 μm. (B) Quantification of cortical enrichment of myosin-II under load. Data is represented as mean ± SD for n≥20 cells analysed for each case, one-way ANOVA, p<0.0005. (C) Time course showing that myosin-II is rapidly recruited to the cell cortex under load. Data given as mean ± SEM, n=10 cells, one-way ANOVA, p<0.005. Cortical enrichment is calculated by measuring the ratio of fluorescence intensity of membrane and cytoplasm around the cell. Ax2 cells expressing myosin-II-GFP (GFP-MhcA) and chemotaxing to cyclic-AMP under 0.5% agarose gels were used throughout. In time courses, load is applied at t=0.

The spatial distribution of myosin-II can be used as a polarity marker and quantified by Fourier analysis (Supplementary figure S3 and Materials and Methods). The distribution of myosin changes from 0.2 ± 0.04 AU under no load condition to 0.1 ± 0.05 AU under 400 Pa load (Supplementary figure S4A). Upon loading, myosin distribution becomes more uniform, with the transition occurring on a similar time scale as recruitment to the cortex (Supplementary figure S4B). This also confirms the visual impression that cells lose polarity when load is applied. An increase in cortical accumulation of myosin is also seen under higher stiffness of agarose gels (Figure 4B, Supplementary figure S4C). However, in these cells myosin-II accumulates preferentially at the rear, unlike the more uniform recruitment in squashed cells.

These results show that load causes persistent myosin-II recruitment to the cell cortex, where it likely increases contractility, so pressurizing the cytoplasm to favour blebbing. They also show that blebbing is not a mere physical response to load but should necessarily involve a biochemical feedback that is triggered by mechanical forces acting on cells.

### Calcium signalling may mediate the response to load

Since cells respond rapidly and sensitively to uni-axial pressure, we hypothesized that a dedicated mechano-sensing system is responsible. One way for cells to sense mechanical force is through an influx of calcium into the cytoplasm mediated by stretch-operated channels. We therefore tested whether the response to load depends on external calcium by using a nominally calcium-free medium, with 200 μM EGTA included to chelate any traces remaining. In this medium, cells continue to move and produce a basal number of blebs (though reduced) but the increase caused by load was virtually abolished (Figure 5A, movie S8). Instead, the cells move predominantly with pseudopods, which constituted more than 90% of the projections produced under a steady-state load of 400 Pa (Figure 5B). The number of blebs produced by wild-type cells treated with EGTA only increased from 0.4 ± 0.3 blebs/cell/min to 0.9 ± 0.1 blebs/cell/min upon application of load, much less than the response without EDTA from 2.1± 0.1 blebs/cell/min to 9.1 ± 0.1 blebs/cell/min (Figure 5B).

**Figure 5.**
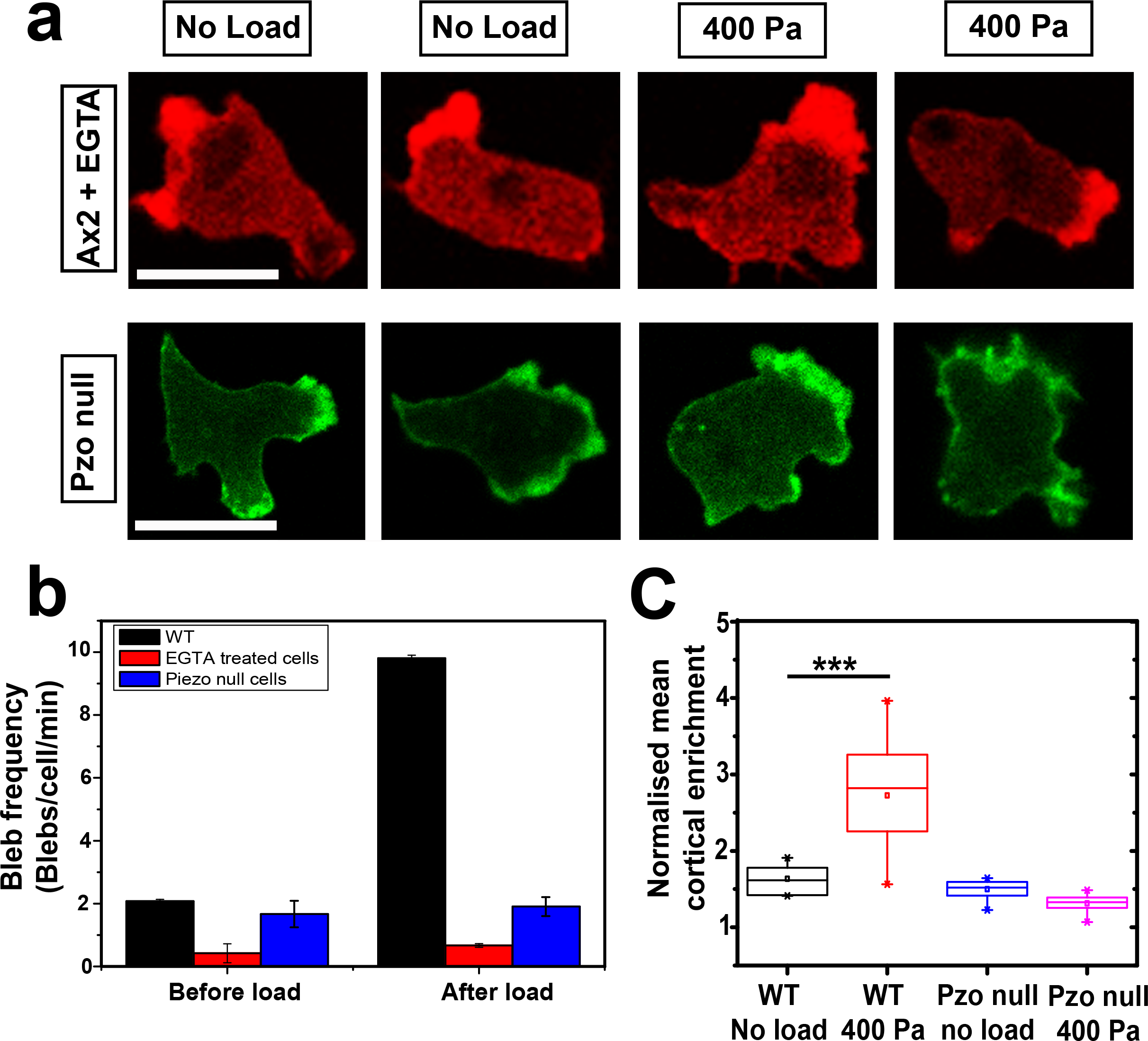
Extracellular calcium and the Piezo stretch-operated channel are required for cells to respond to load. (A). Illustration of typical responses to load of cells either in calcium-free medium, or lacking the Piezo channel (PzoA^−^ cells, strain HM1812). Compare to Ax2 controls in Figures 1C and 2A. Scale bar = 10 μm. (B) Quantification of the blebbing response to load of cells either in calcium-free medium, or lacking the Piezo channel. Data is mean ± SD for n≥15 cells tracked before and after applying load in each case. Cells, either Ax2 parental or Piezo null mutant (pzoA^−^ cells; strain HM1812), were incubated under agarose made with the standard KK2 buffer containing 100 μM calcium or this buffer lacking calcium and supplemented with 200 μM EGTA. A load of 400 Pa was applied as indicated. (C) Quantification of the cortical enrichment of myosin-II in PzoA^−^ cells under a load of 400 Pa. The cortical enrichment of RFP-myosin II is calculated by measuring the ratio of fluorescence intensity of membrane and cytoplasm around the cell. Cells are chemotaxing to cyclic-AMP under 0.5% agarose gels The data is mean ± SD for n≥20 cells analysed for each case, p<0.0005 for wild-type cells and p>0.5 for Piezo null cells, Mann-Whitney test and one-way ANOVA.

To test whether the imposition of load causes an increase in cytoplasmic calcium, we used cells expressing the YC2.60 FRET reporter to measure cytosolic calcium ^48^. The results show that there is an immediate, though modest, increase in the normalised FRET ratio when load is applied to cells: it increases from an average value of 1.0 before load to a maximal of 1.6 shortly after the application of load, returning to baseline of 1.1 in about 20-25 seconds (Figure 6, top panel). The response is clear but much smaller than to a saturating dose of 4 μM cyclic-AMP or 4 μM ionomycin (Supplementary figure S5A). In the presence of EGTA, the normalised FRET ratio hovered around 0.8 with no appreciable increase when load was applied (Figure 6, middle panel), indicating that the increase in cytosolic calcium depends on an influx through the plasma membrane.

**Figure 6:**
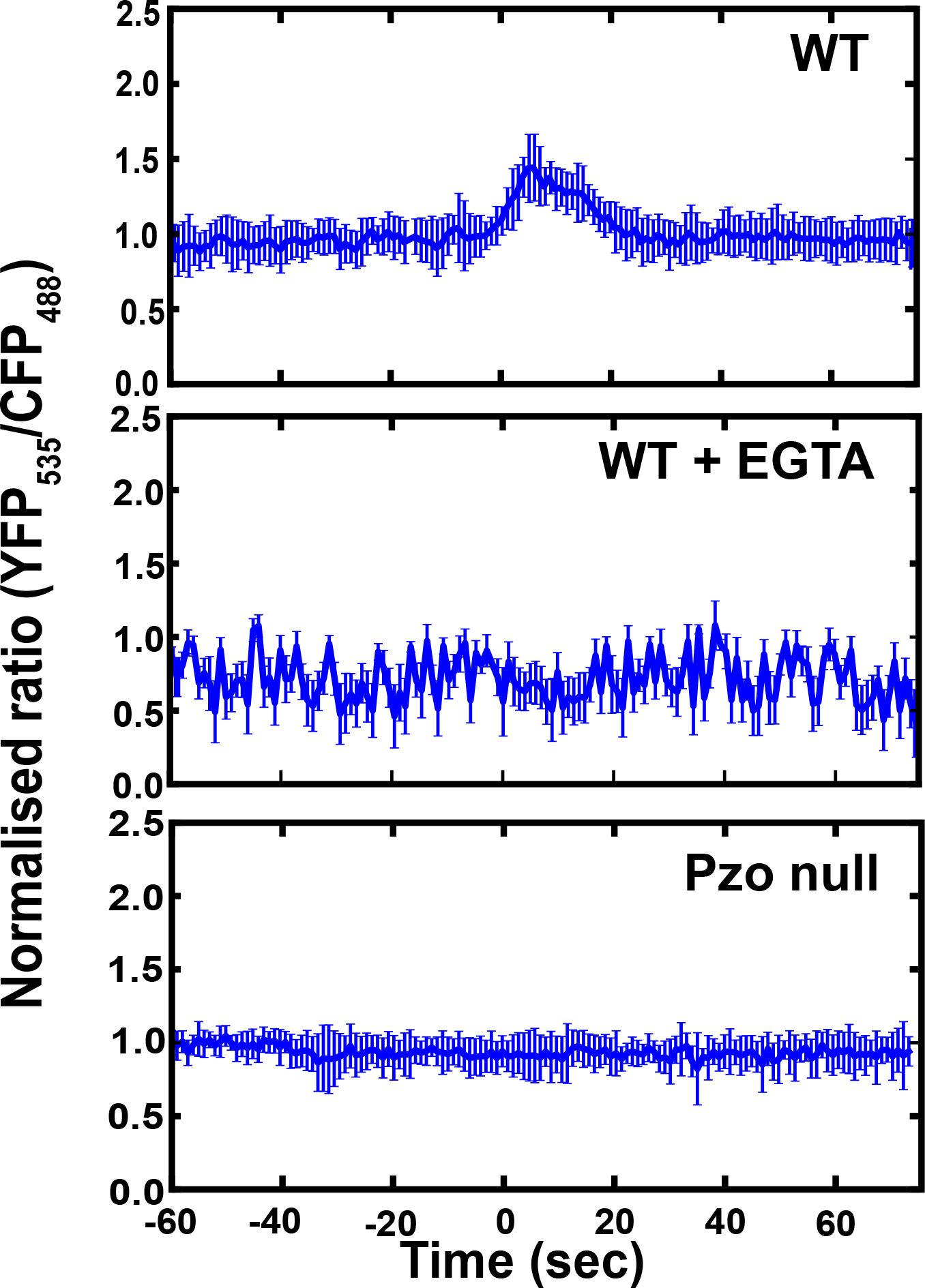
Applying load to cells produces a transient increase in cytoplasmic calcium that depends on extracellular calcium and the Piezo stretch-activated channel. Changes in cytoplasmic calcium were detected by microscopy using the Cameleon FRET-based sensor. The normalised ratio of YFP 535nm/CFP485nm indicates the cytosolic calcium concentration. Aggregation-competent cells under 0.5% agarose were subjected to a load of 400 Pa as indicated (n = 15 cells).

These results therefore suggest that the load applied to cells is sensed by an influx of calcium into the cytoplasm, mediated by an unknown mechano-sensitive channel in the plasma membrane.

### The Piezo stretch-operated channel is required for sensing load

Only a limited number of potential stretch-operated channels are recognizable in the *Dictyostelium* genome ^49^. We screened null mutants in these for defects in responding to load. An existing triple mutant of the mechano-sensing channel MscS and two Trp channels (one a mucolipin homologue, MclN; the other, TrpP, responsive to ATP) ^48^ and a double mutant of IplA (a homologue of the IP3 receptor required for the calcium response to chemoattractants), ^50^ and TrpP channel, both showed a normal response to load, making essentially wild-type numbers of blebs (Supplementary figure S6, movie S9, S10,S11 & S12).

In addition to these channels, the *Dictyostelium* genome encodes a single homologue of the Piezo stretch operated channel (DDB_G0282801 at dictyBase; we designate the gene as *pzoA*). We created knock-out mutants in this gene (Supplementary figure S7), which grew close to normally in shaken suspension in HL5 liquid medium (mean generation times: 8.59 ± 0.11 hr for the Ax2 parent and 9.18 ± 0.12 and 9.56 ± 0.13 for the HM1812 and HM1813 *pzoA*^−^ mutant strains; SEM; n=3).

It is immediately apparent that the Piezo mutant behaves differently under load from its parent, continuing to move with pseudopods instead of blebs (Figure 5A and B). To confirm this, mutant and parent were marked with different coloured fluorescent F-actin reporters, mixed and subjected to a load of 400 Pa: the parent blebs copiously, as expected, but the mutant barely at all (Supplementary figure S8A, movie S13). Instead the mutant continues to move with actin-driven pseudopods. A second Piezo mutant also behaved in a similar way (not shown). Quantification shows that the mutant produces a basal level of blebbing without load, but unlike its parent, load causes little if any increase (Figure 5B, Supplementary figure S8B,E). This is true even for a load of 1600 Pa (Supplementary figure S8B, movie S14). Similarly, inducing the mutant to move under stiffer agarose gives only a small increase in blebbing over basal (Supplementary figure S8C,D), whereas blebbing in the parent increases to around 90% of projections under 2% agarose (Young’s modulus 75 kPa) (Supplementary figure S8D). Cytosolic calcium levels in Piezo null cells were not perceptibly stimulated by load (Figure 6, bottom panel). Although the traces are quite noisy, the normalised FRET ratio fluctuated around 0.8 before the application of load and it did not show any appreciable change upon application of a load of 400 Pa.

Piezo null cells still produce basal levels of blebs, suggesting that blebbing itself does not intrinsically depend on Piezo. To test this, we stimulated cells with cyclic-AMP, which is detected by a G-protein coupled receptor and triggers a burst of blebbing about 20 seconds after acute stimulation ^39^. In this case, cyclic-AMP caused copious blebbing in both wild-type and Piezo mutant cells (Supplementary figure S5B, movie S15). Thus the ability of Piezo null cells to bleb remains intact, but it can no longer be stimulated by mechanical means.

The failure of load to stimulate blebbing in Piezo mutants suggests that they may be defective in myosin-II mobilization. Figure 5C and Supplementary figure S9A,B show that load does not cause myosin-II to be recruited to the cortex of Piezo null cells (movie S16). Additionally, their polarity, as measured by the distribution of myosin-II along the cortex, also does not change when load is applied (Supplementary figure S9C,D).

## Discussion

In their natural habitat of soil and leaf litter *Dictyostelium* cells have to navigate through complex 3D environments as single cells and then adapt to moving within tissues during their multicellular development ^51,52^. In laboratory conditions they show admirable flexibility, being able to move with actin-driven pseudopods under buffer and pressure-driven blebs under an elastic overlay ^41,53,54^. Using a cell squasher ^42^, we show that uniaxial pressure alone is sufficient to rapidly induce *Dictyostelium* cells to switch from moving with pseudopods to blebs. The switch is rapid, occurring in 10 seconds or less, and requires only small pressures of around 100 Pa.

Blebs form when the cell membrane detaches from the underlying cortex and is driven outwards by fluid pressure. Our current biophysical understanding of blebbing involves a balance between the internal hydrostatic pressure of the cell, which favours blebbing, and adhesion between the cell membrane and cortex, which opposes it. Tension in the cell membrane favours or opposes blebbing according to its local curvature ^35,53,54^. The contractility of the cell cortex causes a mechanical tension that is in practice much larger than the tension of the cell membrane bilayer, and is largely responsible for pressurising the cytoplasm according to the Laplace equation. This relates the difference between internal and external pressures to the curvature of the cell interface and cortical tension (Figure 7A). For a cortical tension T of 0.5 mN/m ^55^ and a spherical cell of approximate radius r = 5 μm, the cytosol’s hydrostatic pressure with respect to the buffer is calculated to about P_0_ = 200 Pa.

**Figure 7:**
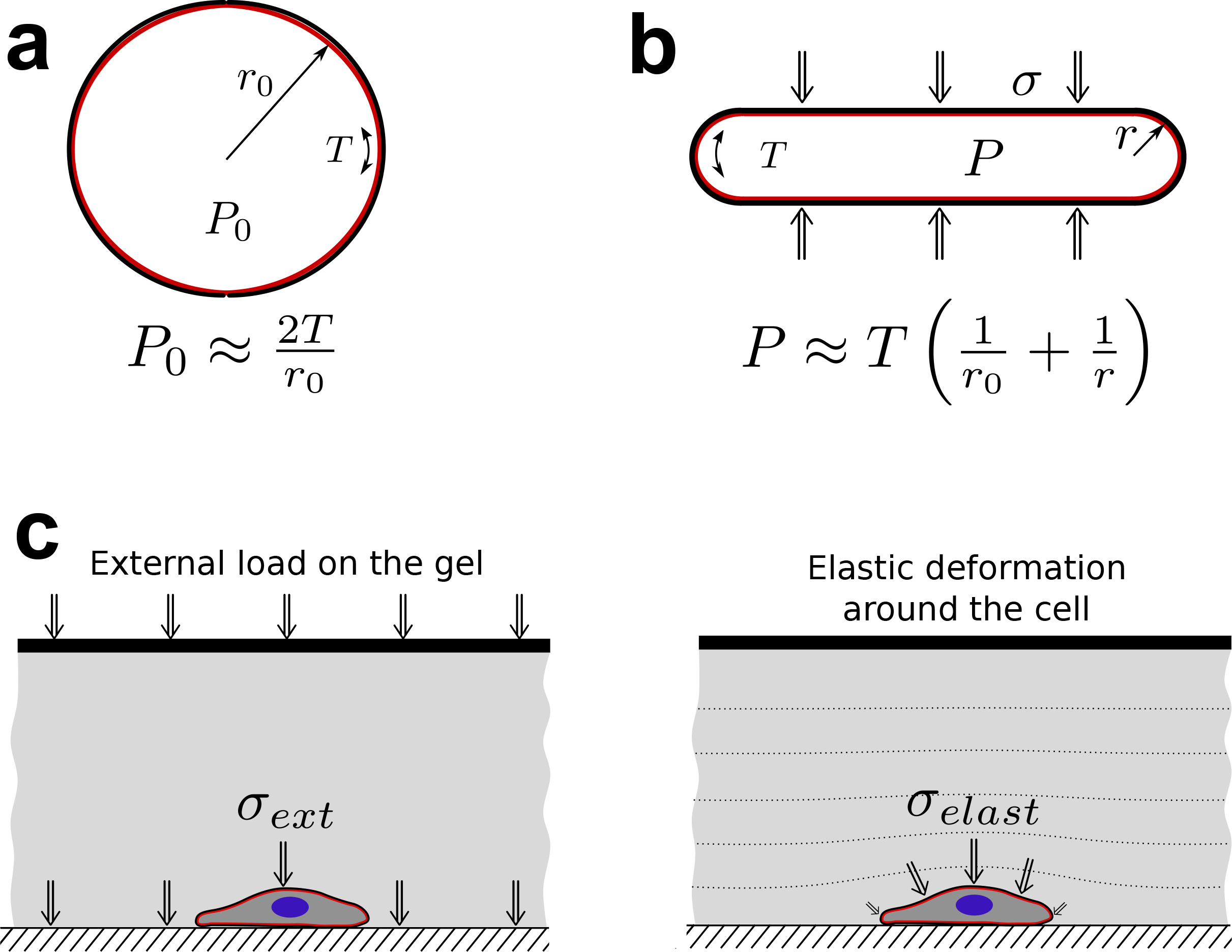
Biophysical representation of the change in cell geometry and pressure balance upon application of external load. (A) Hydrostatic pressure, P_0_, in the cytosol of a spherical cell with a radius r_0_ and cortical tension T. These quantities are linked through the Laplace equation. (B) Application of external load on a spherical cell leads to its flattening and change to a pancake shape. (C) Illustration of how external loading and elastic deformations of the gel contribute to the uniaxial pressure on a cell.

We find that compressing *Dictyostelium* cells with a uniaxial pressure of the same order (a few 100 Pa) as the calculated cytosolic pressure is sufficient to flatten them and halve their height. The resultant increase in cytosolic pressure due to squashing can also be calculated from the Laplace equation. Approximating the squashed cell as a pancake with its smaller radius of curvature r = 2.5 μm at the periphery (Figure 7B) and assuming an unchanged cortical tension, yields an increase in cytosolic pressure of the order of 100 Pa. To balance forces at the top and bottom interfaces, where the curvature is negligible, this pressure must match the externally applied uniaxial stress σ squashing the cell, and we indeed find that an applied load of the order of 100 Pa is required to halve the height of the cells. Thus, this simple mechanobiological picture accounts for the effect of uniaxial loading on the geometry of the cells.

Extending this mechanobiological model, it might be proposed that blebbing is triggered by uniaxial stress because it causes an increase in cytosolic pressure and that this is sufficient to break the linkage between the plasma membrane and cortex. This interpretation is qualitatively consistent with the observed effect of gel stiffness. Under high stiffness 2% agarose, the pressure induced by the gel deformation in the absence of any load is great enough to flatten the cells to the same proportion as a 400 Pa load, and induces a similar effect on blebbing (Figure 7C). From a mechanical point of view, higher load or higher stiffness both result in the same effective confinement of cells, which, combined with their cortical tension, causes an increase in hydrostatic pressure and a qualitatively similar effect on cell migration. This phenomenology also agrees with the observation of blebbing in many mammalian cells confined under a gap of fixed stiffness ^9^, and is therefore likely to be generic.

In the context of this purely mechanical explanation for load-induced blebbing, our finding of a mechanosensory system to detect changes in the mechanical load is surprising. Our results show that the blebbing response to pressure is almost abolished in mutants of the Piezo stretch-operated channel, although these cells remain contractile and fully capable of blebbing when stimulated with a chemoattractant. This demonstrates that squashing the cell alone is insufficient to cause blebbing: some function of the Piezo channel is also required. This establishes that the transition to blebbing migration is under biochemical control.

Piezo channels are inherently mechanosensitive and when opened by membrane tension allow various cations into the cell, including Ca^2+^ ^28^. As well as requiring Piezo, the blebbing response to uniaxial load depends on extra-cellular Ca^2+^ and is associated with a transient increase in cytosolic Ca^2+^. We therefore propose that it is mediated by an increase in cytosolic calcium due to the opening of Piezo channels in the plasma membrane. It is not clear how this transient Ca^2+^ signal produces a persistent change in cell behaviour that lasts as long as load is applied. Possibly cytosolic Ca^2+^ acts as a switch that is reversed by some other means when the pressure is removed, or that local increases in cytosolic calcium persist in the sub-membranous region of the cytoplasm, but are too small to detect by our methods.

The link from Piezo channel activation to blebbing remains to be fully defined. Myosin-II is essential for blebbing, and its recruitment to the cell cortex under load could increase cortical contractility and hence cytosolic pressure, thus favouring blebbing. Myosin-II recruitment is also stimulated by chemoattractants, but this occurs in the presence of EGTA and in the IplA mutant, where there is no detectable increase in cytosolic calcium ^56–58^, and neither myosin heavy chain kinase nor myosin light chain kinase appear to be directly regulated by calcium ^59^. Thus, the link between a calcium influx mediated by Piezo and increased myosin-II driven contractility is likely to be indirect.

Squashing under agarose causes a considerable re-organization of cells, some of which may also be relevant to blebbing. Loss of paxillin-containing adhesions suggests reduced adhesion of the cell to the substratum and is reminiscent of the behaviour of other cell types, which also become less adhesive when moving in a ‘pressure-driven’ mode ^6^. The loss of 25% of cell volume caused by pressure is surprising, given that freely moving *Dictyostelium* cells can radically change in shape and surface area but usually keep their volume nearly constant ^43^. The loss of volume may therefore be an active response of the cell, which given the extreme crowding of the cytoplasm, could have profound effects on properties such as viscosity ^60^ and affect processes including actin polymerization and de-polymerization ^61^. Similar changes in cell volume also occur when glioma cells invade narrow spaces ^62^.

Many cells, like *Dictyostelium* amoebae, change how they move according to their physical environment. Immune cells do it by squeezing through narrow pores of the blood vessels deforming their nucleus in the process ^63^. Many cancer cell types switch to blebbing under confinement ^5,8,9,16,18,35,64^. We suggest that at least some of these behavioural changes involve sensing pressure through Piezo channels.

## Online Materials and Methods

### Cell culture and Reporters

*Dictyostelium discoideum*, strain Ax2 (Kay laboratory strain; DBS0235521 at http://dictybase.org/) was grown at 22°C on *Klebsiella aerogenes* lawns on SM agar plates (Kay,1987) or axenically in HL5 medium (Formedium). It was transformed with an F-actin reporter, ABD-GFP consisting of the F-actin binding domain of ABP-120 (residues 9-248) fused to GFP driven by the actin15 promoter ^65^(strain HM2040). The expression of ABD-GFP was maintained under the selection of 10 ug/ml of G418 antibiotic.

The GFP-MhcA reporter for myosin-II, was created from an extra-chromosomal GFP expression vector (pDM1407) and pJET-MhcA cloning vector ^66^ and an mCherry-MhcA vector was created similarly using an mCherry expression vector (pDM1208). A fluorescent reporter for Paxillin was created in a similar manner using pDM1407 and a cloning vector for paxillin (pDM722) to create GFP-paxillin.

The piezo gene homologue (DDB0282801) in *Dictyostelium* was knocked out using the vector pDT43 (Supplementary figure S7A). A 1.56 kb 5’region of flanking homology was generated by PCR and cloned into pLPBLP as a ApaI/HindIII fragment. Then a 1.32kb 3’ region of flanking homology was generated by PCR and cloned in as a NotI/SacII fragment to generate pDT43. Primer pairs used for PCR were PZKO1/PZKO2(5’) and PZKO3/PZKO4(3’).

PZKO1:5’-TTTGGGCCCTAAATTTATCCTTTTTCATTGATTTGTTCATCAG-3’
PZKO2:5’-ATTAAAGCTTATGCTAATAATATCGCTGATGCTACACTTGCTG-3’
PZKO3:5’-TAAGCGGCCGCTTGAAGTCTCTGTTGTAGGTTGGAATCC-3’
PZKO4:5’-TAACCGCGGTAATTATTATGATACAATTGCTGATATGGAAGCTGC-3’

Ax2 cells were electroporated with digested pDT43 (digested with ApaI/SacII) and transformants selected with blasticidin (20 μg/ml). Genomic DNA was prepared from wells using GenElute kit (Qiagen). For PCR screening following primers were used: PZ1: 5’- GGAAATAAAAAAATGATAGGATATTTCTTTGTG -3’and PZ2: 5’- TGGTAAAACTGTTTCACAAGTTGCTACTTCC -3’, which bound outside the disruption cassette. PzoA KO cells gave a product at 4.42 kb and were found in 2/14 wells. These were re-cloned and assessed by PCR (Supplementary figure S7B).

### Live-cell microscopy and cell squashing experiments

All microscopy was with cells in KK2 buffer (16.5 mM KH_2_PO_4_, 3.8mM K_2_HPO_4_, 2 mM MgSO_4_, and 0.1 mM CaCl_2_). Cells were brought to aggregation-competence by starving freshly harvested cells in KK2 at 2×10^7^ cells/ml with shaking at 180 rpm. After 1 hour cells were pulsed with 90 nM final cyclic-AMP, every 6 minutes for another 4.5 hours. Aggregation-competence was confirmed morphologically by the formation of small clumps. These cells were then imaged in round glass-bottom dishes (35 mm dish with a 10 mm glass bottom, MatTek corporation) using an inverted laser-scanning confocal microscope (Zeiss 780 or 710) with a 63x/1.4 NA oil-immersion objective. The images were collected using Zen2010 software (Zeiss) and processed using Fiji or Image J. A modified version of under-agarose assay was used, as described previously ^41,42,67^. A thin layer of SeaKem GTG agarose (Lonza biochemicals) in KK2 (different concentrations to prepare gels of different stiffnesses chiefly 0.5%, 0.75%, 1% and 2% w/v) of height 2 mm was poured in a preheated round glass-bottom dish (MatTek).

Two parallel rectangular troughs of sizes 4 mm by 8 mm and 1 mm by 5 mm were cut in the gel, once it solidified. A gradient of cyclic-AMP was formed in the chamber by adding 5 μM cyclic-AMP to the larger well and leaving it for about 30 minutes. Once the gradient formed, 2×10^5^ cells/ml were added to the smaller trough and allowed to migrate. The load was applied using the cell squasher, as described previously ^42^, once the cells had migrated under the gel (about 40 minutes). Briefly, the plunger was bought in close contact with the surface of the gel by manually positioning it using the micrometers and subsequently, a command to apply the load was given. A log file recorded the applied load and the position of the plunger.

### Blebbing assay

This assay was adapted as previously described ^39^. Cells were grown to 5×10^6^ cells/ml, starved and pulsed with cyclic-AMP for about 4.5 hours to make them aggregation competent. 2×10^5^ aggregation competent cells/ml in 200 μl KK2 buffer were allowed to attach to a glass-bottom dish for 20 minutes, after which they were imaged by confocal microscope. 16 μl of 50 μm cyclic-AMP was added to the buffer while recording cells at 2fps.

### Image analysis

Blebs and pseudopodia were identified by their morphological characteristics from the movies. Additionally, they were confirmed using kymographs where blebs could be seen as fast projections and devoid of actin polymerization at the leading edge whereas pseudopodia were identified with their characteristic slow actin dynamics (Figure 1B).

In the movies where load was applied dynamically, blebs were scored at their first occurrence and the total number of blebs was binned in 1 second or 1 minute intervals.

Speed of cells was calculated by automated tracking using QUIMP plug-in in Image J ^68^. The centroid of cells was tracked at each frame of the movie to calculate the total distance travelled by the cell. It was then divided by total time to calculate the speed of individual cell.

Cell height was measured by reconstruction of the z-stacks (taken at 0.4 μm increments) using Imaris (Bitplane). Z-axis elongation which occurs due to mismatch in the refractive index ^69^ was corrected by comparing the true and apparent height of a fluorescent bead of a known height (9.7 μm diameter Fluospheres; Molecular probes). The stacks were corrected by dividing them by 1.97 μm (giving a true z-axis increment of 0.203 μm). Statistical analysis of cell height was done using One Way ANOVA and Tukey’s means comparison test in Origin software (Originlabs). Cell volume was calculated from the reconstructed stacks by calculating the volume of individual voxels occupied by the cells. Surface area was measured by calculating the total area of voxels on the outer surface. Fluorescence intensity was measured as the total florescence intensity of all relevant voxels. The distribution and localization of myosin was analysed using a MATLAB plug-in kindly provided by Douwe Veltman and described in ^70^. A typical example of the analysis is shown in Supplementary figure S3. The plug-in tracks the membrane of the cell by identifying points of maximum intensity at the cell edge. It then draws equidistant normals of 3 pixels in length at intervals along the cell boundary, and calculates the points of maximum intensity along these and their coordinates. The mean fluorescence intensity of the background is then subtracted from the membrane and cytosol respectively. Each maxima on the membrane is mapped onto a corresponding local cytosolic point to compute the ratio between membrane and cytoplasm to give normalized fluorescence intensity values (Supplementary figure S3). The distribution of the fluorescent signal is also quantified by measuring its periodicity. Consider the example of the cells shown in Supplementary figure S3, the top cell (Supplementary figure S3A) has signal localized in restricted regions of the cell while in the bottom cell (Supplementary figure S3B), the signal is localized homogeneously throughout the cell. The distribution can be described by a periodic function, and the Fourier transform of the signal defined as:

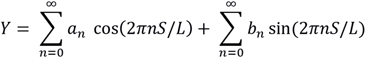

with the ratio of two terms calculated as:

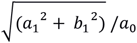

where S: position around cell contour L: perimeter

The first term of the series represents the mean value of the function while the second one is the trigonometric function. The ratio of the first two terms of the series is taken which evaluates the difference in the peaks of the described signal. The more heterogeneous the signal, the larger is the gap between the peaks and as a result, larger is the value of the ratio. In polarized cells the signal is strongly localized to only part of the cell and hence the intensity peaks are further apart, thereby giving a higher ratio between the terms. On the other hand, if the myosin is more homogeneously localized in the cell, then the peaks are clustered together and a lower value of the ratio is expected. The robustness of this method was checked using the cells shown in Supplementary figure S3 and indeed the ratio of the first two terms for a polarized cell (Supplementary figure S3A) was 0.22 while that for un-polarized cell (Supplementary figure S3B) was 0.03.

The lifetime of an actin scar left behind by a bleb was measured from a kymograph taken along the line indicated in Supplementary figure S2A; the tracks demarcating the newly formed bleb (i_bleb_), membrane (i_memb_) and cytoplasm (i_cyto_) are indicated (Supplementary figure S2A). A python script was written to calculate the intensity along the indicated tracks. In order to minimize the error due to fluctuation of fluorescence intensity with position, average intensity values along a line width of 3 pixels was taken. The intensity of scar is then defined as:

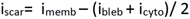

The intensity of the scar is fitted with a sigmoid curve to obtain its half-life. The sigmoid curve used for fitting is:

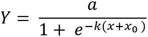

and critical time obtained from the fit is given by

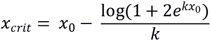

## Acknowledgments

The authors would like to thank the workshops at Department of Engineering and MRC-LMB, Nick Barry for help with microscopy and Radu Tanasa for help in python scripts and Peggy Paschke and Douwe Veltman for help with cloning and generation of expression vectors used in this work. Guillaume Charras, Matthieu Piel and Melda Tozlugolu for the discussions. This work was supported by Dr. Manmohan Singh Scholarship from St. John’s College to N. Srivastava, Medical Research council core funding MC_U105115237 to R.R. Kay and BBSRC grants BB/K018175/1, BB/P003184/1 to A.J. Kabla.

## Supplementary figure legends

**Supplementary figure 1:**
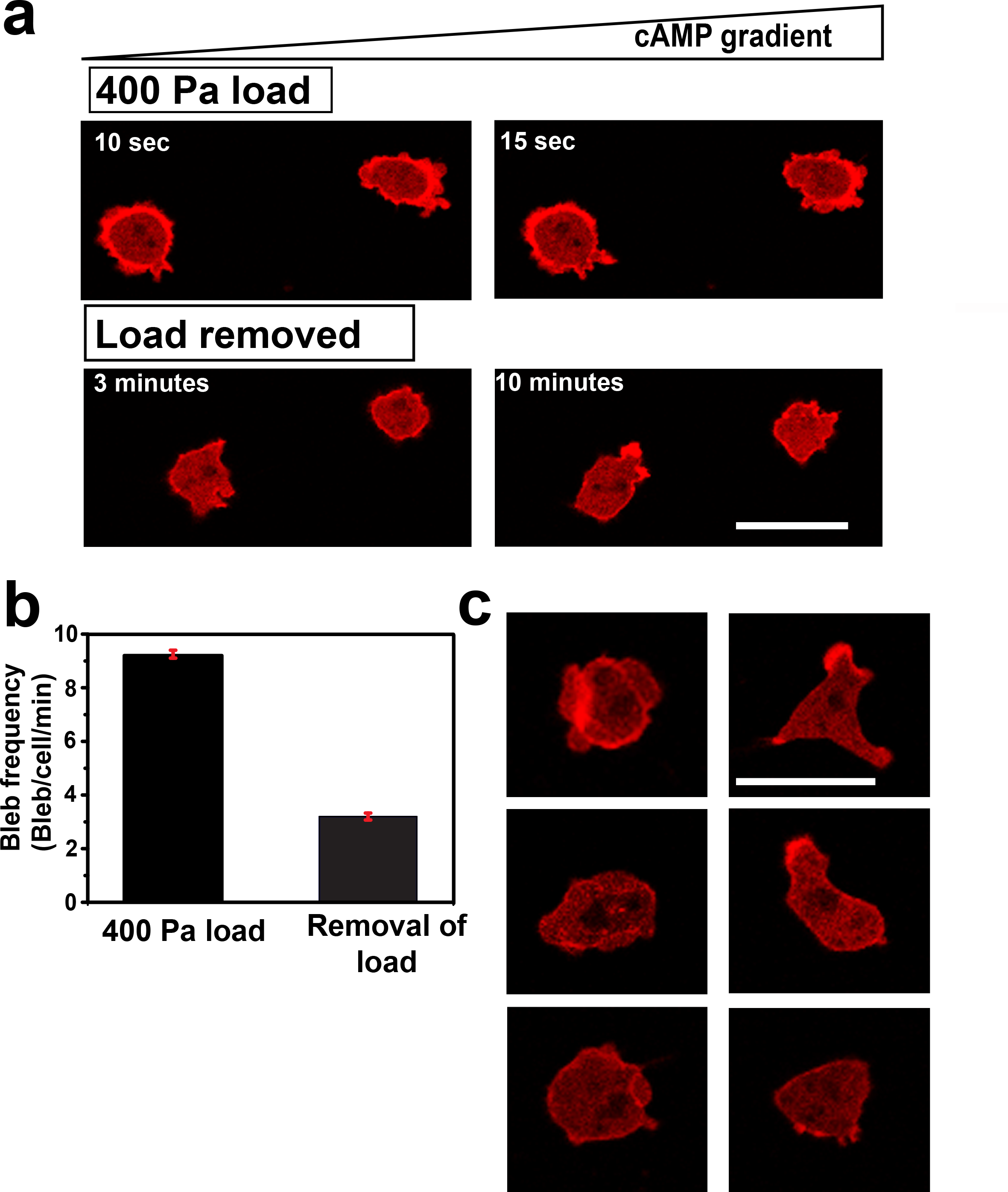
The switch to bleb-driven motility caused by uniaxial load is reversible. (A) Recovery of cells upon removal of external load. Cells revert from blebbing to forming pseudopodia when load is removed. A load of 400 Pa was applied on cells underneath 0.5% agarose gels and removed at t = 0.; Scale bar = 10 μm. (B) Quantification of blebs produced by cells under load and after its removal (after ~10 min). Data is mean ± SD for n≥15 cells tracked before and after the load in each case, p<0.0005 and one-way ANOVA. (C) Montage of cells after removal of load. Wild-type Ax2 cells were compressed by uniaxial 400 Pa and consequently switch to blebbing. Upon removal of load, these cells switch back to forming pseudopodia and adopt a polarised shape after about 10 min; scale bar = 10 μm.

**Supplementary figure 2:**
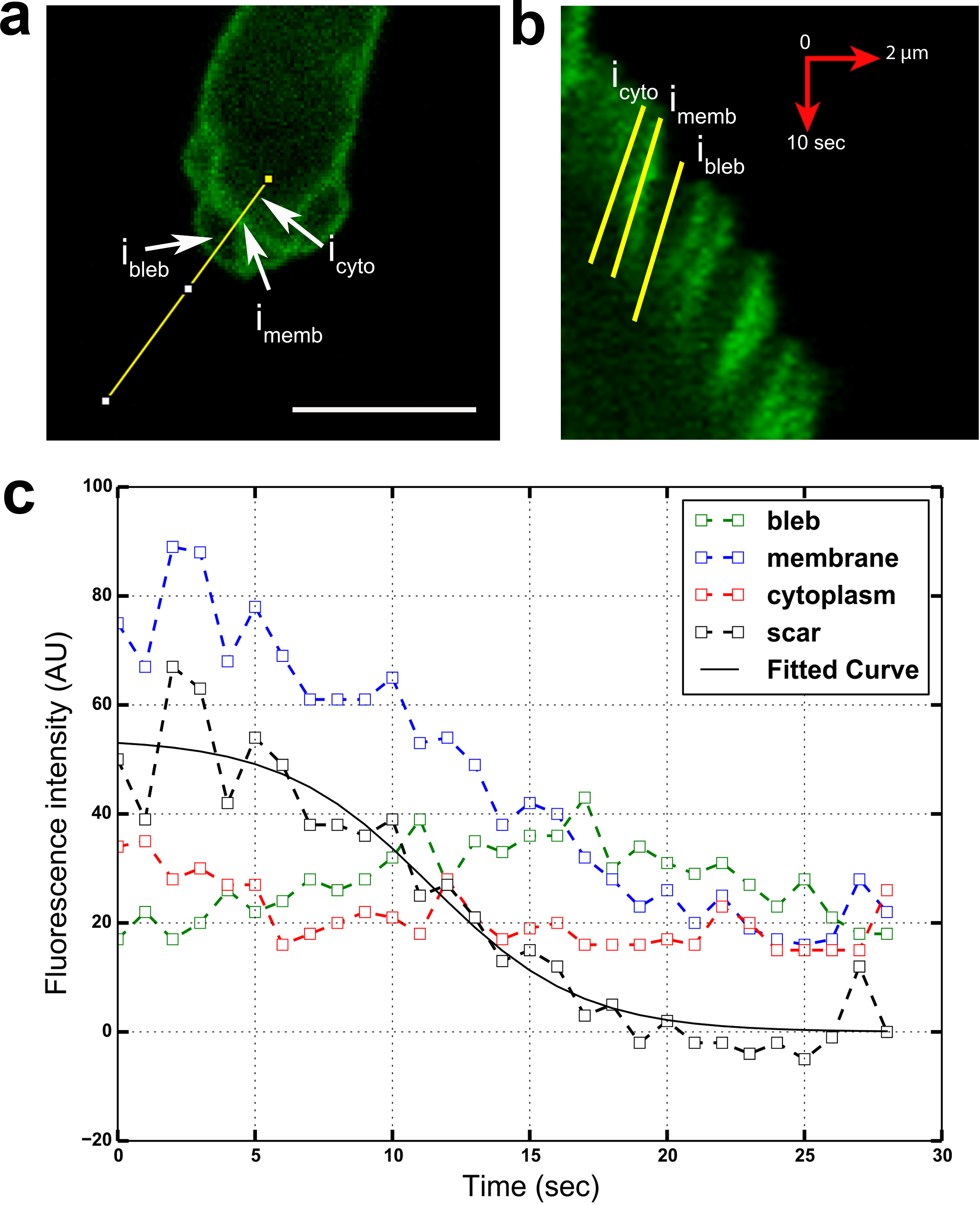
Analysis of the actin scar left behind when a bleb is projected. (A) A cell blebbing under a load of 400 Pa is shown with three distinct regions around a bleb marked as i_bleb_, i_memb_ and i_cyto_ representing the newly formed bleb, the scar left behind by the bleb and cytoplasm of the cell, respectively; Scale bar = 10 μm. (B) Kymograph derived from the line indicated through the bleb shows its progression. Average fluorescence intensity for membrane, cytoplasm and bleb are measured along the paths indicated. (C) Plots of the average fluorescence intensities for membrane, cytoplasm and bleb. Average fluorescence intensity calculated for the actin scar is fitted with a sigmoid function to obtain the half-life.

**Supplementary figure 3:**
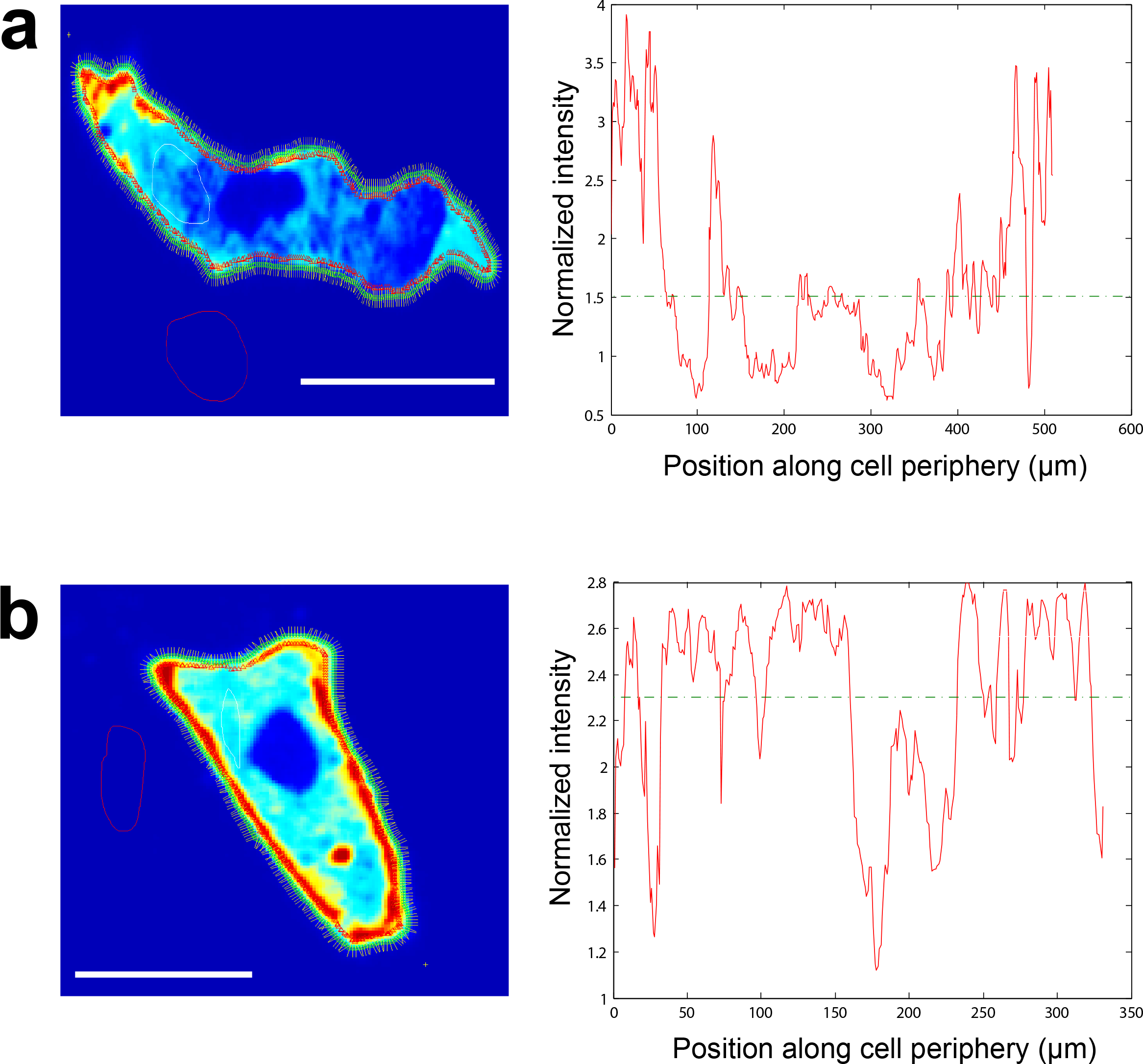
Analysis of the distribution of myosin-II using Fourier analysis. A representative (A) polarized cell and an (B) less polarized cell. The graph indicates the variation in the fluorescence intensity of GFP-MhcA signal at the cell periphery, with a dashed line indicating the average intensity of the measured signal. After Fourier analysis of these intensity values, the ratio of the first and the second term defines the distribution of the intensity signal and thus, the polarity of the cell (see Materials and Methods).

**Supplementary figure 4:**
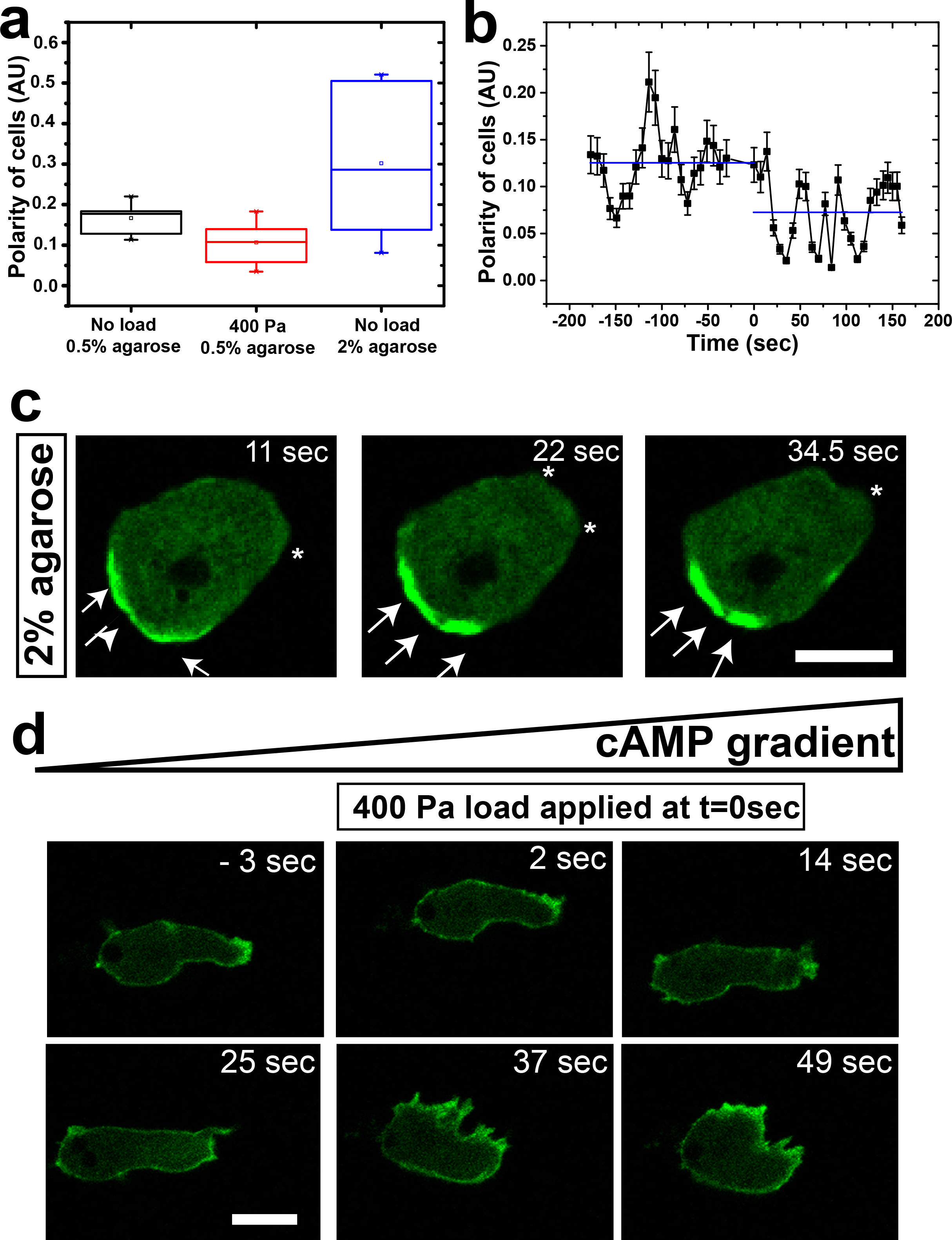
Uniaxial loading causes changes in myosin-II distribution. (A) Time course of changes in cortical GFP-MhcA polarity under load. Data is mean ± SEM for 10 cells, each tracked as load is applied starting at 0 sec; one-way ANOVA, p<0.005. Fourier transform of membrane fluorescence intensity values is used to quantify the polarity of cells (see supplementary Figure 4 for details). (B) Quantification of GFP-MhcA cortical enrichment under load. Data is represented as mean ± SD for n≥20 cells analysed for each case, one-way ANOVA, p<0.005. (C) Dynamics of myosin-II enrichment in cells expressing GFP-MhcA migrating under stiff 2% agarose gel; Scale bar = 10 μm. (D) Myosin-II null cells fail to bleb under uniaxial load, arguing that blebbing is an active process requiring contractility generated by myosin-II. Uniaxial load of 400 Pa was applied to MhcA null cells at t=0. The cells are expressing the F-actin reporter, ABD-120 GFP, and are under 0.5% agarose gel; scale bar: 10 μm.

**Supplementary figure 5:**
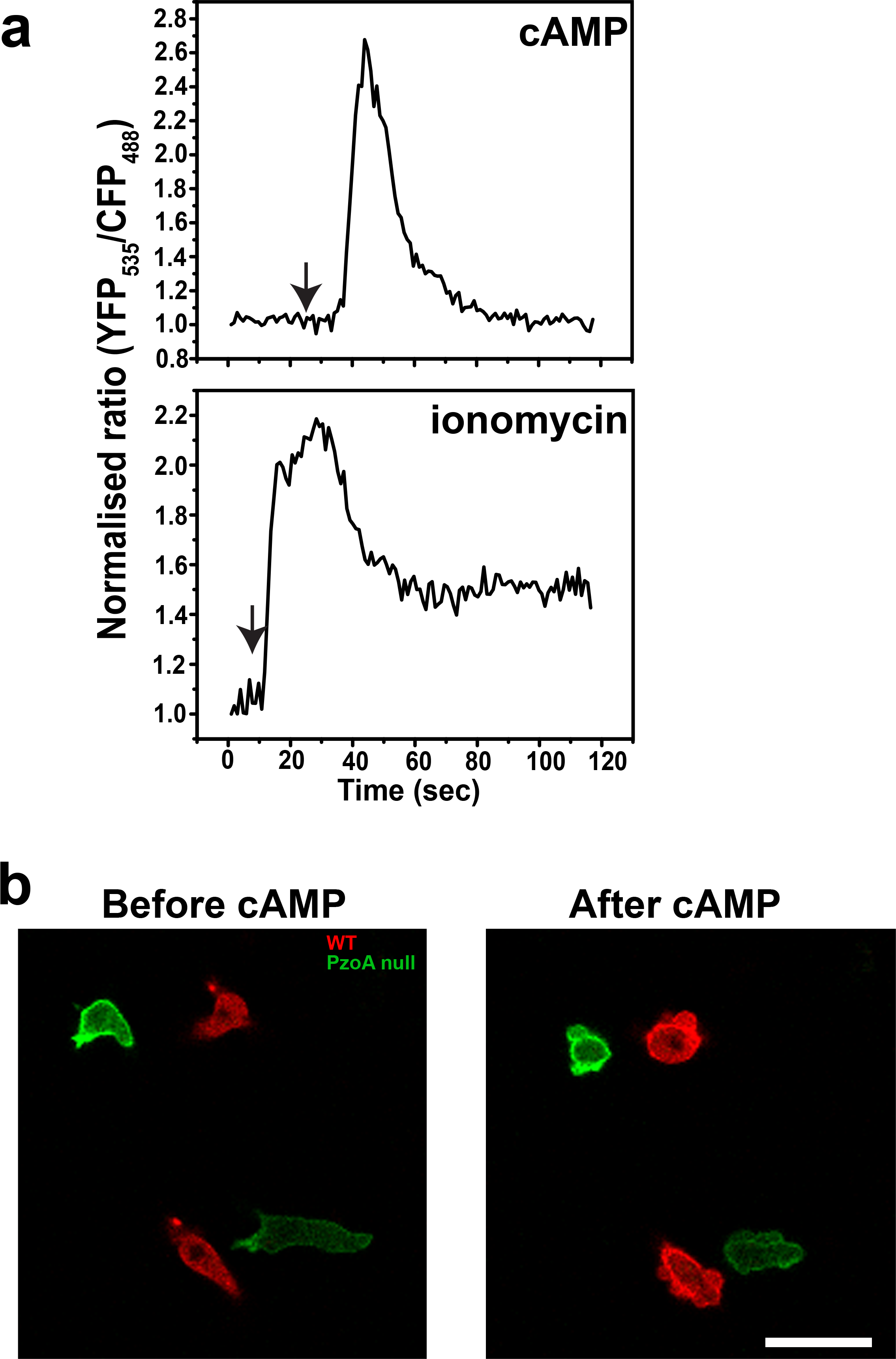
Effect of cyclic-AMP and ionomycin on cytoplasmic calcium levels and the blebbing response of wild-type and Piezo null cells to cyclic-AMP. (A) Cytoplasmic calcium levels are elevated following the addition of cyclic-AMP or ionomycin to wild-type Ax2 cells. Cytosolic calcium levels were measured using the Cameleon FRET-based sensor, with the normalised ratio of YFP 535nm/CFP485nm indicating the cytosolic calcium concentration (n = 20 cells). (B) Cyclic-AMP shock causes transient blebbing in a both wild-type cells expressing LifeAct-RFP and PzoA^−^ cells expressing LifeAct-GFP, scale bar: 10 μm.

**Supplementary figure 6:**
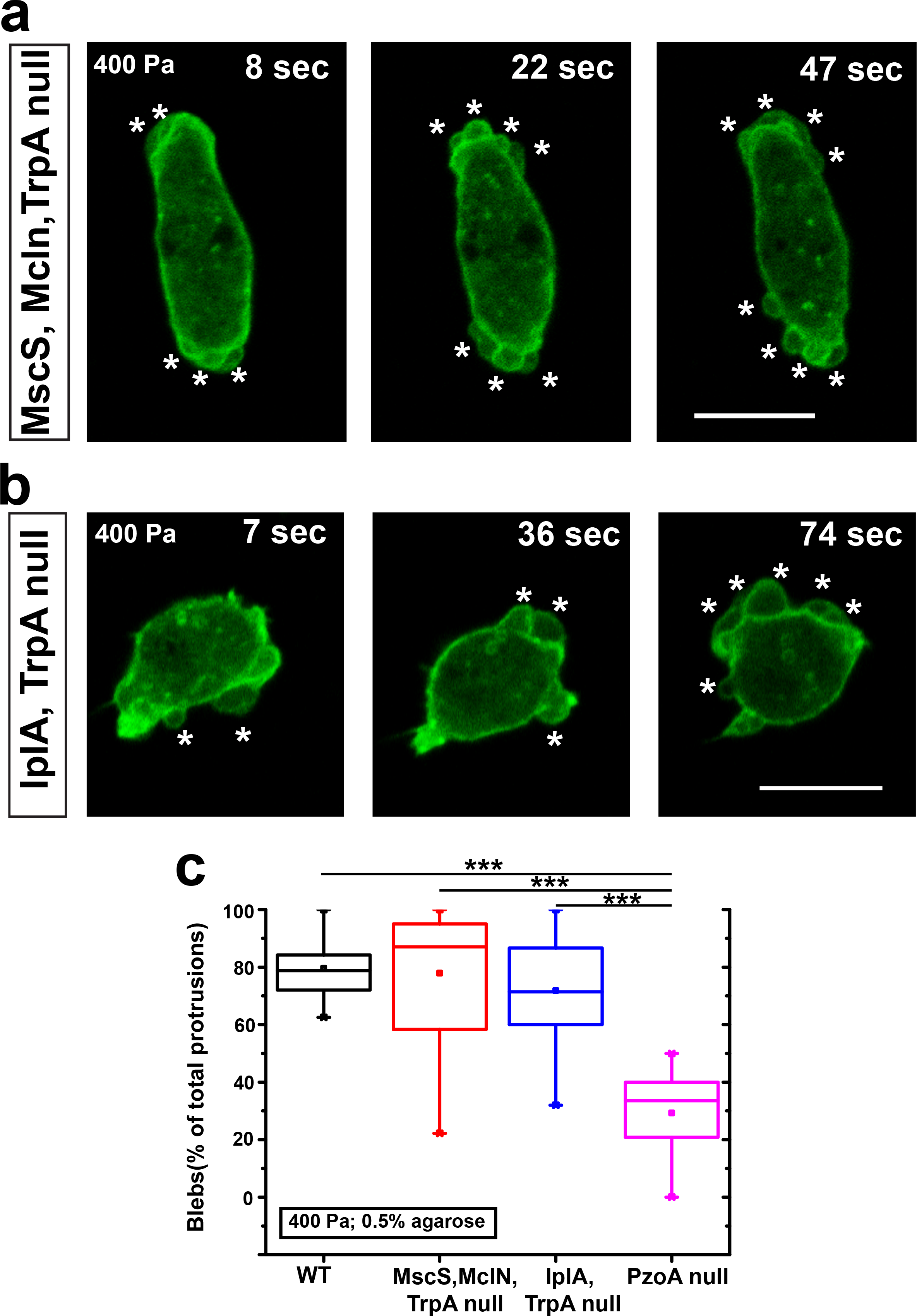
Effect of uniaxial loading on mutant cells lacking potential stretch-sensitive ion channels. (A) Triple mutant of MscS, mucolipin (MclN) and TrpP channel. (B) Double mutant of IplA and TrpP channel. (C) Quantification of blebbing after application of uniaxial load to strains mutant for candidate mechanosensitive ion channels and wild-type Ax2 cells. Scale bars are 10 μm.

**Supplementary figure 7:**
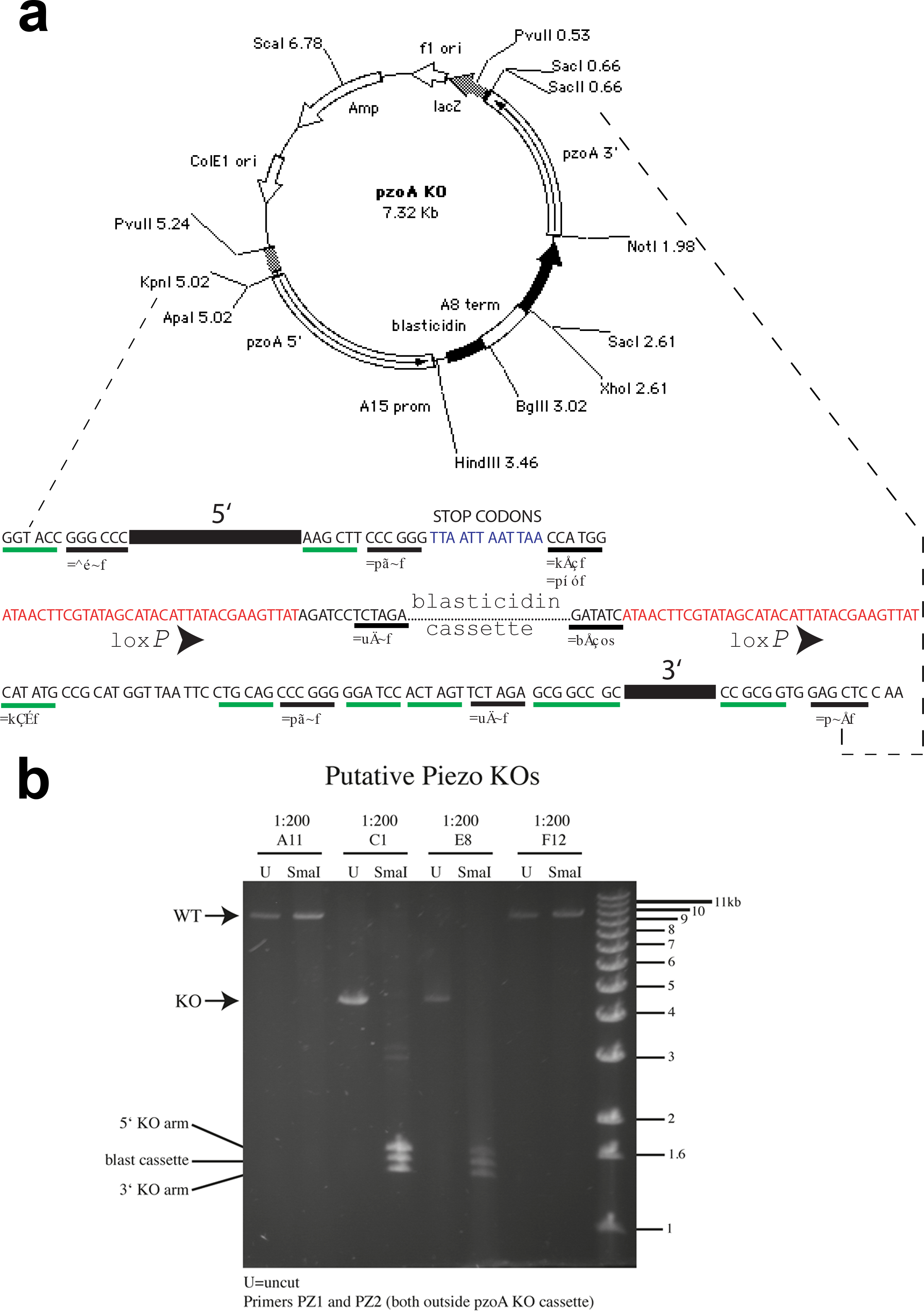
Creation of PzoA knock-out strains. (A) PzoA knock-out strains were created using the PzoA knock-out plasmid, pDT43 (plasmid design by Macvector software). (B) A diagnostic PCR showing two positive clones, which yield a band at around 4.5 kB, which is sensitive to digestion with SmaI. This liberates the 5’ and 3’ homology arms together with the blasticidin cassette, while in wild-type cells no product was obtained.

**Supplementary figure 8:**
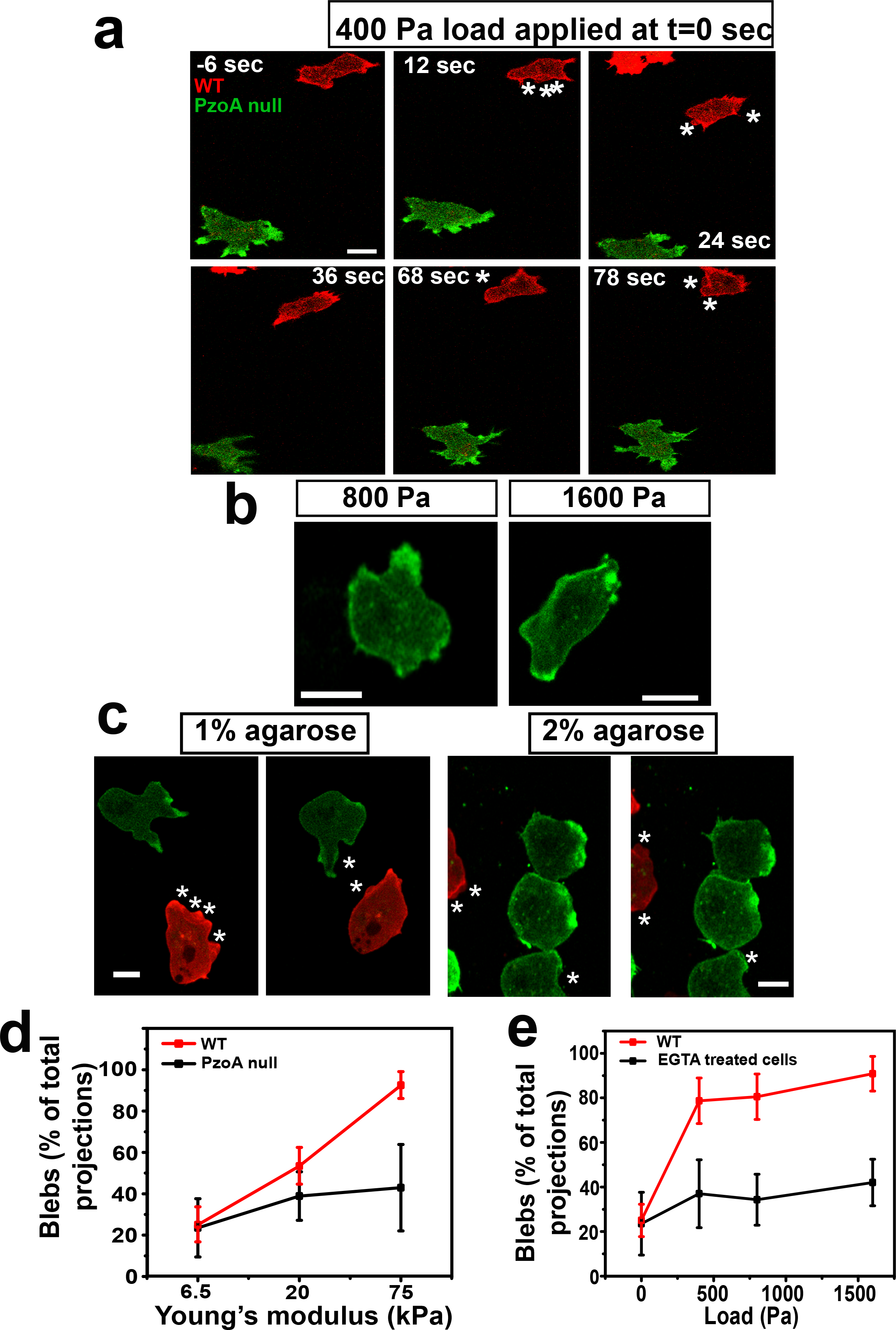
Piezo is required for sensing compressive load. (A) Piezo null cells (PzoA^−^) do not respond to uniaxial load by blebbing. Load is applied to a mixture of PzoA^−^ cells expressing LifeAct-GFP and wild-type cells expressing LifeAct-mRFP under 0.5% agarose. PzoA^−^ cells continue to migrate with pseudopods when the load is applied but wild-type cells form blebs instead (indicated by *) (Scale bar:10 μm). (B) Greater uniaxial loading does not cause blebbing in PzoA^−^ cells. PzoA^−^ cells were subjected to 800 Pa and 1600 Pa load but continued to migrate with pseudopods; Scale bar = 10 μm. (C) Increasing the stiffness of agarose gels does not cause a switch to blebbing in PzoA^−^ cells. A mixture of PzoA^−^ cells expressing LifeAct-GFP and wild-type cells expressing LifeAct-mRFP were induced to chemotax under stiffer 1% and 2% agarose gels. PzoA^−^ cells moved with fewer blebs compared to wild-type cells; Scale bar = 10 μm. Quantification of blebs formed by wild-type and PzoA^−^ cells under (D) stiff agarose gels and (E) compressive load. The data is represented as mean ± SD for n ≥ 30 cells for each case.

**Supplementary figure 9:**
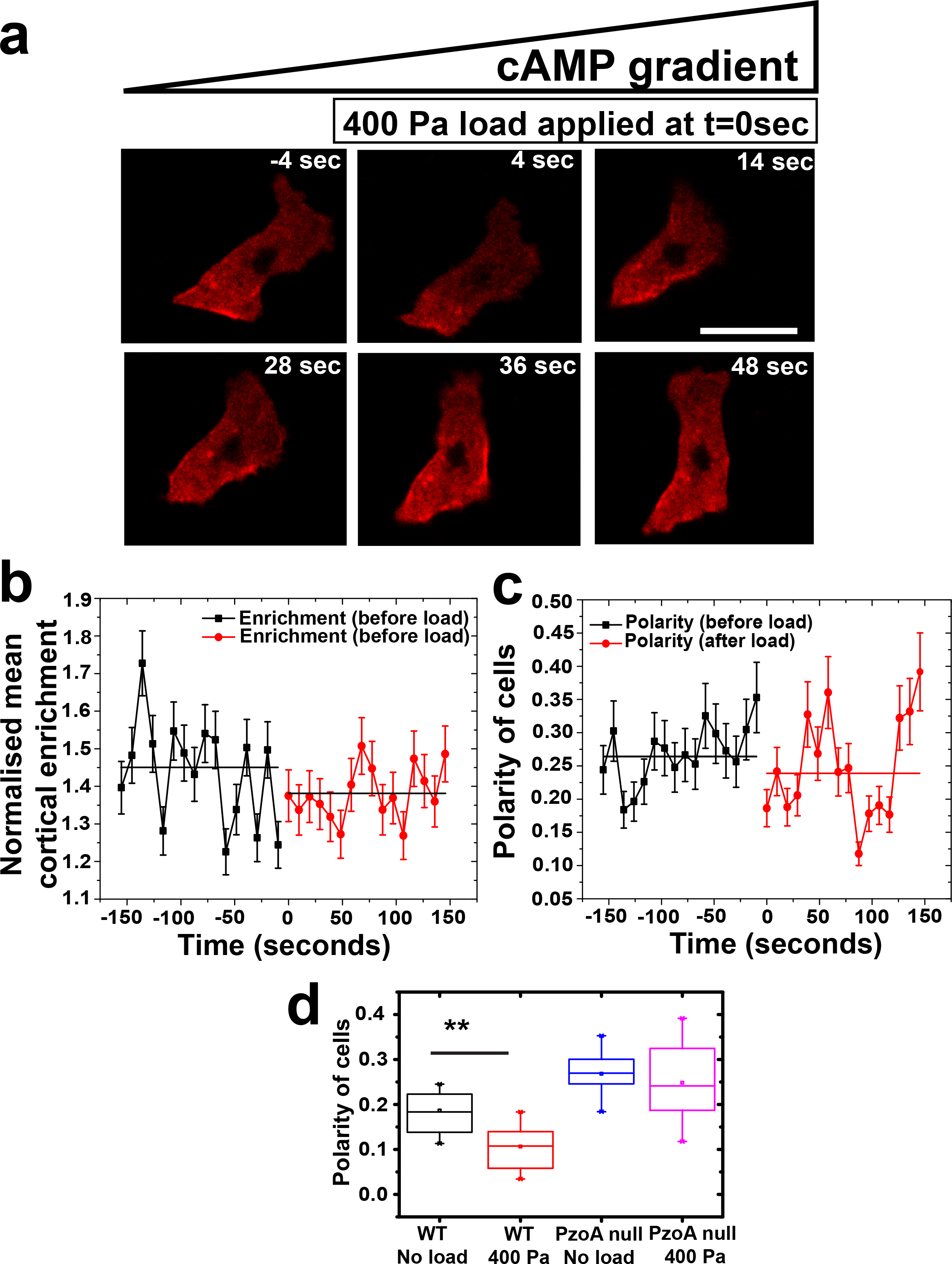
Myosin distribution does not change under compressive load in PzoA^−^ cells. (A) Myosin dynamics in PzoA^−^ expressing RFP-myosin-II (RFP-MhcA) during loading. Load was applied to cells chemotaxing under 0.5% agarose gel. Time is given with respect to the start of loading (t=0); blebs are indicated by * and scale bar =10 μm. (B) Time course of RFP-MhcA recruitment to the cell cortex under load in PzoA^−^ cells. (C) RFP-MhcA distribution under load in PzoA^−^ cells. Time is given with respect to the start of loading (t=0). Data is represented as mean ± SEM, n=10 cells, one-way ANOVA, p>0.5 in both cases. (D) Quantification of the cortical distribution of myosin-II in wild-type and Piezo null cells under a load of 400 Pa. The data is represented as mean ± SD for n≥20 cells analysed for each case, p<0.005 for wild-type cells and p>0.5 for Piezo null cells, Mann-Whitney test and one-way ANOVA.

## Caption for movies

**S1: Cell moving under 0.5% agarose gel.** Aggregation competent Ax2 cell expressing ABD-GFP as a reporter for F-actin, is chemotaxing to cyclic-AMP under 0.5% agarose without external mechanical load applied. Cells move predominantly with a pseudopod and rarely with blebs under these conditions. Taken by a laser scanning confocal microscope (Zeiss 780) at 2 frames per second.

**S2: Cell moving under 100 Pa compressive load and 0.5% agarose gel**. An aggregation competent Ax2 cell expressing ABD-GFP as a reporter for F-actin, is chemotaxing to cyclic-AMP under 0.5% agarose and a compressive load of 100 Pa. The cell migrates with a combination of blebs (indicated by *) and pseudopods. Majority of blebs are formed at the leading edge. Taken by a laser scanning confocal microscope (Zeiss 780) at 2 frames per second about 8-10 minutes after the load is applied.

**S3: Cell moving under 800 Pa compressive load and 0.5% agarose gel**. An aggregation competent Ax2 cell expressing ABD-GFP as a reporter for F-actin, is chemotaxing to cyclic-AMP under 0.5% agarose and a compressive load of 400 Pa. The cell typically migrates with blebs (indicated by *) and infrequently with pseudopods. Taken by a laser scanning confocal microscope (Zeiss 780) at 2 frames per second about 8-10 minutes after the load is applied.

**S4: Cell migrating under 1600 Pa compressive load and 0.5% agarose gel**. An aggregation competent Ax2 cell expressing ABD-GFP as a reporter for F-actin, is chemotaxing to cyclic-AMP under 0.5% agarose and a compressive load of 1600 Pa. The cell typically projects numerous blebs (indicated by *) with a drastic decrease in the migratory speed. The blebs typically form all around the cells with a noticeable change in the cell shape. Taken by a laser scanning confocal microscope (Zeiss 780) at 2 frames per second about 8-10 minutes after the load is applied.

**S5: Instantaneous response of a cell to a 400 Pa.** An aggregation competent Ax2 cell expressing ABD-GFP as a reporter for F-actin is chemotaxing to cyclic-AMP under 0.5% agarose, at first without an external load. It is then suddenly subjected to a compressive load of 400 Pa. The cell initially moves by forming pseudopods under 0.5% agarose but starts to bleb (indicated by *) as soon as the load is applied. Taken by a laser scanning confocal microscope (Zeiss 780) at 2 frames per second.

**S6: Switch to blebbing mode of migration is reversible upon removal of the compressive load.** Aggregation competent Ax2 cells expressing LifeAct-mcherry as a reporter for F-actin and chemotaxing to cyclic-AMP under 0.5% agarose gel and 400 Pa compressive load show blebbing phenotype. Subsequently, the blebbing decreases as the load is removed. The cells revert back to pseudopod-based migration with a distinct polarized shape by about 10 min after removal of load. This clearly shows that the switch to blebbing is reversible. Taken by a laser scanning confocal microscope (Zeiss 780) at 2 frames per second in the beginning of the movie and 1 frame per 2 minutes after the removal of the load.

**S7: Rapid recruitment of myosin to the cell cortex upon application of compressive load on cells.** Aggregation competent Ax2 cells expressing GFP-MhcA as a reporter for myosin-II are coaxed to chemotax towards cyclic-AMP under 0.5% agarose gel. Myosin, which is localized mostly at the rear and the sides of the migrating cell, rapidly relocalizes to the cell cortex in response to 400 Pa compressive load. The temporal dynamics of myosin loacalization is similar to the timescale of protrusion switching from pseudopodia to blebs due to compressive loading. Taken by a laser scanning confocal microscope (Zeiss 780) at 2 frames per second.

**S8: Cells do not switch to bleb driven migration under compressive loading in the absence of extracellular calcium.** Aggregation competent Ax2 cells expressing ABD-GFP as a reporter for F-actin, in calcium-free media to which 200 μM EGTA was added, chemotaxing towards cyclic-AMP under 0.5% agarose gel. The cells predominantly migrate with pseudopods underneath the soft gel and do not switch to bleb-driven migration when a compressive load of 400 Pa is imposed. These cells continue to form pseudopods as they migrate. Taken by a laser scanning confocal microscope (Zeiss 780) at 2 frames per second.

**S9: A triple mutant of mucolipin, TrpP and MscS channels migrating under a compressive load.** An aggregation competent triple mutant of the mechano-sensing channel MscS and two Trp channels (one a mucolipin homologue, MclN; the other, TrpP, responsive to ATP) expressing LifeAct-GFP as a reporter for F-actin, is chemotaxing to cyclic-AMP under 0.5% agarose and a compressive load of 400 Pa. The cell predominantly migrates by forming blebs and very infrequently with pseudopods. Taken by a laser scanning confocal microscope (Zeiss 780) at 2 frames per second about 8-10 minutes after the load is applied.

**S10: Instantaneous response of a triple mutant of mucolipin, TrpP and MscS stretch-activated channels to a compressive load of 400 Pa.** Aggregation competent cells of the triple mutant of the mechano-sensing channel MscS and two Trp channels (one a mucolipin homologue, MclN; the other, TrpP, responsive to ATP) expressing LifeAct-GFP as a reporter for F-actin, is chemotaxing to cyclic-AMP under 0.5% agarose. The cell typically moves by forming pseudopods under the soft gel but rapidly switches to bleb-driven migration as a dynamic load of 400 Pa is imposed upon it. Taken by a laser scanning confocal microscope (Zeiss 780) at 2 frames per second.

**S11: A double mutant of TrpP and IplA stretch-activated channels migrating under a compressive load.** An aggregation competent double mutant of the mechano-sensing channels: Trp channel (TrpP, responsive to ATP) and IplA (a homologue of the IP3 receptor required for the calcium response to chemoattractants), expressing LifeAct-GFP as a reporter for F-actin, is migrating under 0.5% agarose and a compressive load of 400 Pa, by predominantly forming blebs. Taken by a laser scanning confocal microscope (Zeiss 780) at 2 frames per second about 8-10 minutes after the load is applied.

**S12: Instantaneous response of a double mutant of TrpP and IplA stretch-activated channels to a compressive load of 400 Pa.** An aggregation competent double mutant of the mechano-sensing channels: Trp channel (TrpP, responsive to ATP) and IplA (a homologue of the IP3 receptor required for the calcium response to chemoattractants), expressing LifeAct-GFP as a reporter for F-actin, is chemotaxing to cyclic-AMP under 0.5% agarose. The cell initially moves by forming pseudopods under the soft gel but switches to a bleb-driven migration as a load of 400 Pa is imposed upon it. The response to load by this mutant is similar to that of its WT parent. Taken by a laser scanning confocal microscope (Zeiss 780) at 2 frames per second.

**S13: Instantaneous response of cells lacking the Piezo channel to a compressive load of 400 Pa.** A mixture of aggregation-competent Ax2 cells expressing LifeAct-RFP and *pzoA*- null cells expressing LifeAct-GFP is chemotaxing towards cyclic-AMP underneath a 0.5% agarose overlay. A compressive load of 400 Pa applied to the cells causes a switch from pseudopodia to bleb-driven motility in WT cells (blebs are indicated by *), whereas *pzoA^−^* mutants are seemingly unresponsive, and continue to form pseudopods. Taken by a laser scanning confocal microscope (Zeiss 780) at 2 frames per second.

**S14: Migration of Piezo null cell under 0.5% agarose gel to which a 1600 Pa compressive load is applied**. The cell forms both pseudopods and blebs, unlike its wild-type parent, which exclusively forms blebs under the same conditions. Taken by a laser scanning confocal microscope (Zeiss 780) at 2 frames per second about 8-10 minutes after the load is applied.

**S15: Stimulation of blebbing by cyclic-AMP in wild-type and Piezo null cells.** A mixture of aggregation competent Ax2 cells, expressing LifeAct-RFP, and Piezo null cells, expressing LifeAct-GFP are stimulated with a saturating dose of 4 μM cyclic-AMP. This causes both wild-type and Piezo mutants to transiently round up and copiously bleb, showing that Piezo null cells retain the ability to bleb in response to chemotactic stimulation but not mechanical load. Cell in in KK2 buffer imaged by laser scanning confocal microscopy (Zeiss 780) at 2 frames per second.

**S16: The effect of compressive load on myosin-II distribution in Piezo null mutant cells.** Aggregation competent Piezo null mutant cells expressing RFP-MhcA as a reporter for myosin-II chemotaxing towards cyclic-AMP under 0.5% agarose gel. Similar to the parent Ax2 cells, myosin-II in Piezo mutants localizes to the rear and side of the cells, but unlike the parent, the application of a load of 400 Pa does not cause recruitment of myosin-II to the cortex. Taken by laser scanning confocal microscopy (Zeiss 780) at 2 frames per second.

